# An eco-physiological model of forest photosynthesis and transpiration under combined nitrogen and water limitation

**DOI:** 10.1101/2023.10.11.561680

**Authors:** Peter Fransson, Hyungwoo Lim, Peng Zhao, Pantana Tor-ngern, Matthias Peichl, Hjalmar Laudon, Nils Henriksson, Torgny Näsholm, Oskar Franklin

## Abstract

Although the separate effects of water and nitrogen (N) limitations on forest growth are well known, the question of how to predict their combined effects remains a challenge for modeling of climate change impacts on forests. Here, we address this challenge by developing a new eco-physiological model that accounts for plasticity in stomatal conductance and leaf N concentration. Based on an optimality principle, our model determines stomatal conductance and leaf N concentration by balancing carbon uptake maximization, hydraulic risk and cost of maintaining photosynthetic capacity. We demonstrate the accuracy of the model predictions by comparing them against gross primary production estimates from eddy covariance flux measurements and sap-flow scaled canopy transpiration in a long-term fertilized and an unfertilized Scots pine (*Pinus sylvestris* L.) forest in northern Sweden. The model also explains the response to N fertilization as a consequence of (i) reduced carbon cost of N uptake and (ii) increased leaf area per hydraulic conductance. The results suggest that leaves optimally coordinate N concentration and stomatal conductance both on short (weekly) time scales in response to weather conditions and on longer time scales in response to soil water and N availabilities.

## Introduction

Human-made increases in atmospheric carbon dioxide (CO_2_) concentration have led to rising temperature, and more drought events (IPCC 2014), which have major impacts on gross primary production (GPP) and forest growth. On the one hand, one might expect higher temperature to positively affect biomechanical processes of photosynthesis (Sage and Kubien 2007) and hence growth. On the other hand, more drought events increase the risk of hydraulic failure and higher mortality (McDowell et al. 2008, Ryan 2011). In addition, tree growth is limited by other factors, such as nitrogen (N) availability, which is particularly important in boreal forests in the northern latitudes (Tamm 1991, Binkley and Högberg 2016, Högberg et al. 2017). Thus, interactive effects of water and temperatures on photosynthesis and growth are further influenced by plant nutrition and soil N accessibility, but it is not yet clear how to best incorporate them in process-based models.

Process-based physiological models are well-suited for assessing the response of photosynthesis to different climate drivers. One such model is the well-established Farquhar and von Caemmerer model of leaf photosynthesis (Farquhar et al. 1980, Farquhar and von Caemmerer 1982). To account for resource limitation of photosynthesis, this model can be complemented with models of stomatal conductance and photosynthetic capacity, which is linked to leaf N concentration. Several semi-empirical models have been proposed to model the response of stomata, such as the Ball and Berry model (Ball et al. 1987), where the stomatal conductance (*g*_*s*_) is linearly related to the quantity *AH*_*r*_/*C*_*a*_. Here, *A* is the carbon assimilation rate, *H*_*r*_ is the relative humidity, and *C*_*a*_ is the ambient CO_2_ concentration at the leaf surface.

A limitation of the empirical models is that they can be safely applied only within the range of environmental conditions and observations for which they were developed, and thus may not be accurate under novel conditions or climate change. To overcome this limitation, adaptive models based on optimization principles have been developed. These models assume that the responses of *g*_*s*_ and other plant variables to environmental variations are regulated by an optimal trade-off between carbon gain and cost. In the case of *g*_*s*_, the apparent cost is the loss of water through transpiration. Based on this premise Cowan & Farquhar (1977) proposed the optimal water use efficiency hypothesis (WUEH) where the carbon gain was *A* and the cost was assumed proportional to leaf transpiration (*E*), i.e. net gain is *A* − *E*/*λ*, where *λ* is a constant. However, observations have shown that the cost does not merely increase linearly with respect to *E*, but the slope steepens with rising *E*, as a result of increased absolute water potential and thus bringing the vascular system closer to xylem cavitation (Wolf et al. 2016). Mathematically, this means that the derivative of the cost function, with respect to *E*, should be an increasing function of *E*, defined, for example, as a concave-up parabola (Wolf et al. 2016, Anderegg et al. 2018) or a sigmoid (Sperry et al. 2017). Following the approach of Sperry et al. (2017), Eller et al. (2018) proposed the SOX model in which the costs is a function of root–canopy hydraulic conductance, *k*_*rc*_. The underlying assumption is that the cost will increase as the *k*_*rc*_ decreases due to the increase in absolute water potential necessary to maintain *E*. A similar assumption is also applied in the model by Sabot et al. (2022).

While the above-mentioned models have only considered the cost associated with water transport, other models have also incorporated the cost of maintaining photosynthetic capacity into the cost term (Friend 1991). Following similar ideas, Prentice et al. (2014) proposed that the cost of maintaining photosynthetic capacity is proportional to *V*_c,max_/*A*, where *V*_c,max_ is the maximum rate of RuBP carboxylation. The drawback of the Prentice et al. (2014) model is similar to that of the Cowan & Farquhar (1977) approach, in that the cost associated with transpiration is proportional to *E*/*A*, thus its slope is not an increasing function of *E* (Sabot et al. 2022). In contrast, a more recent approach by Joshi et al. (2022) and Flo et al. (2023), called the P-hydro model, optimizes both photosynthetic capacity and has a transpiration cost increasing with *E* linked to increasing negative plant water potential assumed to cause hydraulic limitation and damage (Joshi et al. 2022). Also, the recent model by Sabot et al. (2022) optimizes not only the trade-off between hydraulic function and photosynthesis but also optimizes photosynthetic N and its distribution between different components, even accounting for the limited adjustment rates and associated delay of leaf N adjustments over time. However, none of the above-mentioned models explicitly accounts for the effects of varying soil N availability.

In this study, we present a new leaf optimization model which combines the cost of maintaining photosynthetic capacity, inspired by the P-hydro model (Joshi et al. 2022), with the hydraulic cost representation of the SOX model. Thus, we optimize not only stomatal conductance as in the SOX model, but also the leaf N content. Our model also accounts for the difference in time-scale between the regulation of stomatal conductance and leaf N content. This leaf-based optimality model is upscaled to allow calculations of canopy GPP and transpiration (*E*_*c*_). In contrast to existing models of this type (Sabot et al. 2022, Joshi et al. 2022) we include the cost of N uptake in order to account for variation in soil N availability. We test and validate the model against observed GPP from eddy covariance flux measurements and *E*_*c*_ data for a Scots pine (*Pinus sylvestris* L.) forest in northern Sweden, where 13 years of controlled annual fertilization has been administered alongside an untreated reference stand. This setting allows us to test our model with varying soil N availability and variable climate over several years. We show that the model predicts the seasonal pattern of *GPP* and *E*_*c*_ well. It also predicts the differences between control and N-fertilized plots as a consequence of different carbon costs of N uptake and leaf area per sapwood area.

## Theory and model

### Model description

A flowchart of the model is provided in Fig. 1 and detailed descriptions of the sub-models are presented in the succeeding subsections. Our model calculates daily stand-level canopy gross primary production (GPP) and canopy transpiration (*E*_*c*_) based on leaf area index (LAI), canopy height (*H*), and climate data (see Table 1 for a full list of necessary inputs). We assume that physiological response is controlled by two plastic variables: the stomatal conductance (*g*_*s*_) and foliage N mass-based concentration (*N*_*m*,*f*_) of a leaf/needle situated at the top of the canopy. The plastic variables are determined by optimization with respect to a fitness proxy, which represents the net carbon gain per leaf area. Central to the optimization is the instantaneous fitness proxy, *G*, which is calculated as the instantaneous leaf-level carbon assimilation of a leaf/needle situated at the top of the canopy (*A*) subtracted by the cost of maintaining transpiration and photosynthetic capacity (the potential electron transportation of a leaf situated at the top canopy, *J*_max_). The cost of maintaining transpiration reflects drought-related loss of soil-canopy conductance (*k*_sc_) due to xylem embolism, which corresponds to a proportional loss of function of the supported leaf area and its carbon assimilation. The cost of maintaining potential photosynthetic capacity is *N*_r_+*N*_*u*_ related to two contributing processes: (i) respiration of a leaf and its supporting roots and stem tissues (*N*_r_), which is assumed proportional to leaf N content, and (ii) the carbon cost of N uptake (*N*_*u*_), which depends on soil N availability. The calculation procedure of *G* is as follows: first, *A* and *J*_max_ are calculated using a mechanistic physiological model. *A* and *J*_max_ are functions of climate variables (above canopy photosynthetic active radiation, *I*_0_, ambient air temperature, *T*_*a*_, ambient carbon dioxide partial pressure *C*_*a*_, and vapor pressure deficit, (VPD) and the two plastic variables. Next, *k*_sc_ is calculated by the hydraulic model with VPD, soil volumetric water content (*θ*), canopy height (*H*), and *g*_*s*_ as input. Using these calculations in an iterative optimization algorithm the cumulative fitness, i.e., the integral of *G*, over a week is maximized by optimizing *N*_*m*,*f*_ and daily *g*_*s*_ values (two *g*_*s*_ values for each day and one *N*_*m*,*f*_ value for the entire week). Subsequently, daily GPP and *E*_*c*_ are calculated by upscaling the leaf-level values to the stand-level. Here, the stand LAI and the daylight hours (Δ*t*_*g*_) are used as additional input.

**Table 1:** Model input. Value ranges are taken from weather data and modeled size data for the fertilized and control stand at Rosinedal experimental site (2015-2018). The data measurements are from 2015-2018, during the growth period.

| Symbol | Description | Range (units) |
| --- | --- | --- |
| *Climate and environmental input* |  |  |
| $T_a$ | Ambient air temperature | -5.1-30.8 (°C) |
| $I_0$ | Above canopy photosynthetic active radiance | 1.8-63.5 (mol m<sup>-2</sup> day<sup>-1</sup>). These values are estimated from solar radiation data using the conversion rate 1000 W m<sup>-2</sup> ≈ 2300 μmol m<sup>-2</sup> s<sup>-1</sup> (2.2.1.a, Landsberg & Sands, 2010) |
| $C_a$ | Ambient CO<sub>2</sub> partial pressure | 40 (Pa) |
| $VPD$ | Vapor pressure deficit | 45.5-1420 (Pa). Estimated from vapor pressure data and calculated saturated vapor pressure values. Saturated vapor pressure is calculated as a function of $T_a$ (Alduchov and Eskridge 1996) |
| $\theta$ | Soil volumetric water content | 9.9-29.8 (%) (-0.42 - -0.13 MPa soil water potential) for the fertilized stand and 6.7-21.1 (%) (-0.64 - -0.19 MPa soil water potential) for the control stand |
| $\Delta t_g$ | Daylight hours | 10.2 - 20.3 (h). The daylight hours are calculated using eq 17 from Jenkins, (2013) |
| *Stand and tree size input* |  |  |
| $H$ | Canopy height | 19.07-19.87 (m) for the fertilized stand and 20.86-21.36 (m) for the control stand |
| $LAI$ | Leaf area index (projected) | 2.38-2.45 (m<sup>2</sup> m<sup>-2</sup>) for the fertilized stand and 2.21-2.30 (m<sup>2</sup> m<sup>-2</sup>) for the control stand |

**Fig. 1.**
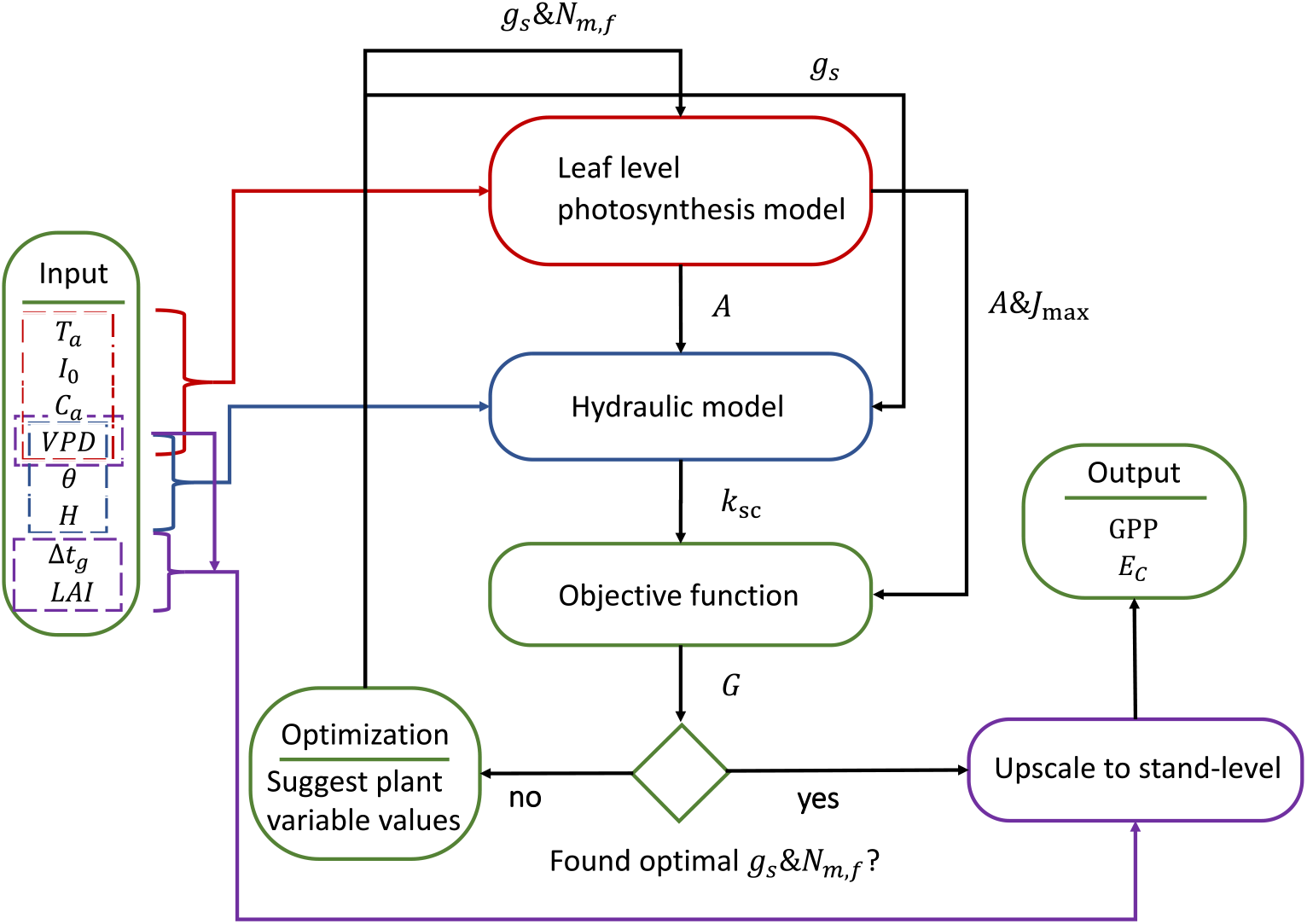
The model is composed of four main modules: Leaf level photosynthesis model (LPM), hydraulic model (HM), objective function (OF), and upscaling module (UM). These modules are accompanied by an optimization routine (OR). To run the model, the following inputs are required: ambient air temperature (T_a_), above canopy photosynthetic active radiance (I_0_), ambient carbon dioxide partial pressure (C_a_), vapor pressure deficit (VPV), soil volumetric water content (θ), daylight hours (Δt_g_), canopy height (H), and stand leaf area index (LAI). OR is used to determine the value of the plant variables, stomatal conductance (g_s_) and mass-based foliage nitrogen concentration (N_m,f_) of a leaf situated at the canopy top, by maximizing the trait performance measure (G). G is determined by OF with soil-canopy conductance (k_sc_), carbon assimilation (A) and potential rate of electron transport (J_max_) as input. A and J_max_ are calculated by LPM with T_a_, I_0_, C_a_, VPV, g_s_, N_m,f_ as inputs. k_sc_ is calculated by HM with VPV, θ, H, and g_s_ as inputs. Once an optimum is found, UM upscales leaf-level values to stand-level and we get the model output: per ground area canopy gross primary production (GPP), and canopy transpiration (E_C_).

### Leaf level photosynthesis model

The leaf level carbon assimilation is calculated in a standard fashion as a balance between the rate of assimilation (carbon demand) and the mass transport of carbon dioxide into the leaf through stomatal and mesophyll conductance (carbon supply). The assimilation rate, A (mol m^-2^ s^-1^) is calculated as the minimum of electron transport limited assimilation rate, *A*_*j*_, and the carboxylation-limited assimilation rate, *A*_*c*_, (Farquhar et al. 1980). We assume co-limitation, i.e., that *A*_*c*_ = *A*_*j*_ (coordination hypothesis, Chen et al., 1993; Maire et al., 2012; Wang et al., 2017; Smith et al., 2019). Further, we assume that the carbon assimilation rate *A* = *A*_*j*_, thus,

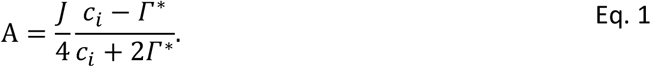

In Eq. 1, *c*_*i*_ (Pa) is the intercellular partial pressure of carbon dioxide, *Γ*^*^ (Pa) is the carbon dioxide compensation point, and *J* (mol m^-2^ s^-1^) is the rate of electron transportation, see Eq. 2.

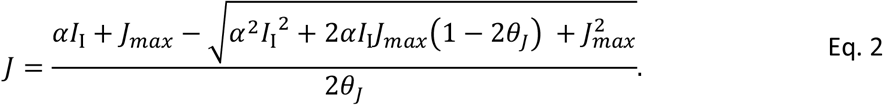

In Eq. 2, *I*_I_ (mol m^-2^ s^-1^) is the irradiance incident on a leaf, *J*_*max*_ (mol m^-2^ s^-1^) is the potential electron transportation, *θ*_*J*_ (-) is a measure of the curvature of the light response curve, and *α* (-) is the quantum yield.

The mass transportation of carbon dioxide into the chloroplast through stomatal and mesophyll conductance is given by Fick’s law:

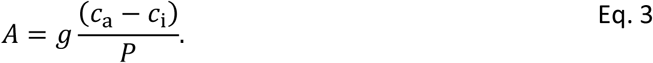

In Eq. 3, *g* (mol m^-2^ s^-1^) is the combined stomatal (*g*_*s*_) and mesophyll (*g*_*m*_) conductance, and *G* is the atmospheric pressure (Pa). We assume that *g*_*m*_ is proportional to *g*_*s*_, thus *g* ≈ *g*_*s*_. Specifically, we assume *g* ≈ 0.42*g*_*s*_ (Wang et al. 2017). Because the demand and the supply equations need to be balanced, we get:

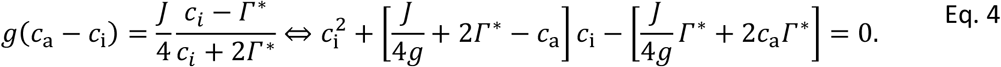

Eq. 4 is a quadratic equation and *c*_i_ is given by the greater of the two roots. Thus, we have an expression for *c*_i_ as a function of *g, I*_I_, and *J*_*max*_, i.e., *c*_i_(*g, I*_I_, *J*_max_). Similarly, we get an expression for the carbon assimilation (Eq. 5) by substituting Eq. 4 into Eq. 3.

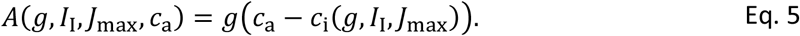

### Temperature dependency of photosynthetic parameters and acclimation to annual temperature cycle

We use the Arrhenius equation and the model from Tarvainen et al. (2018) to estimate the short-term temperature dependency of the photosynthetic parameters. Specifically, the temperature responds of *Γ*^*^, is modelled by using the Arrhenius equation (Landsberg and Sands 2010),

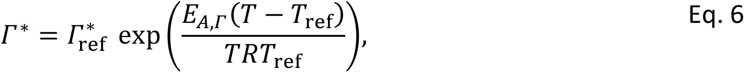

In Eq. 6, *R* = 8.314 (J K-1 mol-1) is the gas constant, *E*_*A*,*Γ*_ (J) is the activation energy of the parameter, *T* is the temperature (K), and 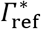 is the parameter value at a reference temperature *T*_ref_ (298 K).

We use Eq. 4 from Tarvainen et al. (2018) to model the short-term temperature responds of *J*_*max*_,

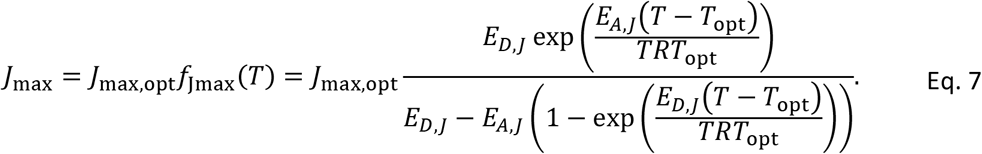

In Eq. 7, *E*_*D*,*J*_ (J) is the deactivation energy, *E*_*A*,*J*_ (J) is the activation energy, and *J*_max_ is the value of *J*_max_ at optimal temperature *T*_opt_ (K).

On a longer timescale, parameters may acclimate to the annual temperature cycle. Specifically, the magnitude of the light response curve follows the trend of the temperature cycle while the shape of the curve remains constant (Hari and Mäkelä 2003, Mäkelä et al. 2004). Mäkelä et al. (2004) enforced this property by assuming that the quantum yield is proportional to *J*_*max*_. For our model, we achieve the same effect by assuming that both *J*_*max*_ and the quantum yield, α, follow the same seasonal cycle, specifically, *α* = *X*_*t*_ *α*_season_ and *J*_max_ = *X*_*t*_ *J*_max,season_(*T*_*a*_, *N*_*a*_). Here, *α*_season_ and *J*_max,season_(*T*_*a*_, *N*_*m*,*f*_) are parameters representing the seasonal apex of *α* and *J*_max_, respectively, and *X*_*t*_ ∈ [0,1] is a variable accounting for the reduction of *α* and *J*_max_ due to the seasonal variation in temperature. Note that *J*_max,season_(*T*_*a*_, *N*_*m*,*f*_) depends on the ambient air temperature, *T*_*a*_, and the nitrogen concentration per leaf mass, *N*_*m*,*f*_. If these equations are substituted into Eq. 2 we get Eq. 8.

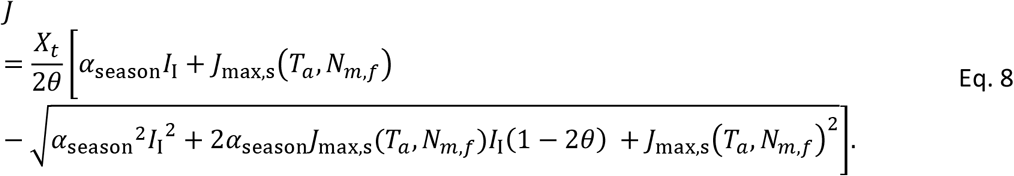

The time-dependent variable *X*_*t*_ (-) is a function of the delayed ambient air temperature, *S*_*t*_ (°C), see Eq. 9.

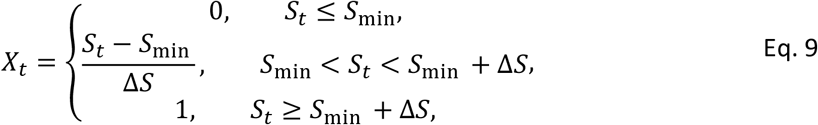

In Eq. 9, *S*_min_ is a parameter representing the minimum threshold for the activation of photosynthesis, and Δ*S* ≥ 0 is a parameter controlling when the photosynthetic capacity reaches its seasonal peak. Thus, *J*_max_ increases linearly with respect to *S*_*t*_ in the temperature range *S*_min_ < *S*_*t*_ < *S*_min_ + Δ*S*. The delayed temperature, *S*_*t*_, is the effective temperature to which the photosynthesis has acclimated to, i.e. the temperature that determines the level of activation of photosynthesis (*X*_*t*_, eq. 9). Because this acclimation takes time *S*_*t*_ lags behind the current temperature, which is modelled using a first order delay dynamics model (Mäkelä et al. 2004, 2008):

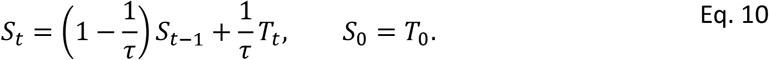

In Eq. 10, *T*_*t*_ is the ambient air temperature at time *t, T*_0_ is an initial temperature of a temperature time series, and *τ* is a parameter controlling the temperature delay; A higher value of *τ* equals a longer delay in the temperature response.

### The effect of nitrogen concentration on the photosynthetic capacity

We assume that the seasonal apex of *J*_max,opt_ is proportional to the per leaf-mass nitrogen concentration, *N*_*m*,*f*_, i.e., *J*_max,opt_(*N*_*m*,*f*_) = *a*_*Jmax*_*N*_*m*,*f*_ (Franklin 2007, Landsberg and Sands 2010). Here, *a*_*Jmax*_ is a proportionality parameter. Thus, *J*_max,season_(*T*_*a*_, *N*_*m*,*f*_) = *J*_max,opt_(*N*_*m*,*f*_) *f*_Jmax_(*T*_*a*_) and *J*_*max*_ = *X*_*t*_ *J*_max,opt_(*N*_*m*,*f*_)*f*_Jmax_(*T*_*a*_).

### Hydraulics model

If *g*_*s*_ and VPD, are given, we can calculate the canopy transpiration per leaf area, *E*:

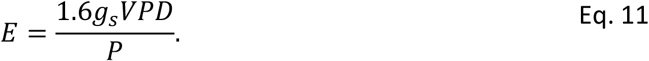

In Eq. 11, *P* is the atmospheric pressure (Pa). We assume that the water flow between root and leaf is in steady-state and negligible non-stomatal water loss, which means that *E* equals the water uptake. We use Darcy’s law to calculate the canopy water potential, *ψ*_*c*_ (MPa), as a function of soil water potential, *ψ*_*s*_, and *E* (Eller et al. 2018):

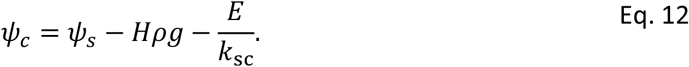

In Eq. 12, *ρ* = 997 (kg m^-3^) is the density of water, *g* = 9.82 (m s^-2^) is the gravitational acceleration, and *k*_*sc*_ (mol m-^2^ leaf s-^1^ MPa-^1^) is the soil-canopy conductance. *k*_sc_ decreases from a potential maximal value, *k*_sc,max_, as water potential, *ψ*, declines according to the vulnerability function, *P*(*ψ*):

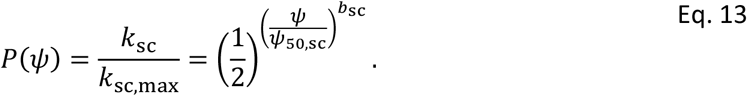

In Eq. 13, *ψ*_50,*sc*_ is the water potential resulting in half of the maximum conductivity, i.e., *P*(*ψ*_50,*rc*_) = 0.5, and *b*_sc_ is a shape parameter controlling how fast *k*_sc_ decreases with the water potential.

We calculate *ψ*_*s*_ from the effective soil saturation, *S*_*e*_ (-), by applying equation 2.19 from Jansson & Karlberg (2011):

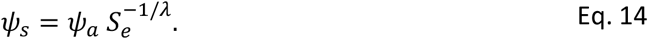

In Eq. 14, *ψ*_*a*_ is the air-entry tension and *λ* (-) is the pore size distribution index of the soil. The effective saturation is a function of soil water content (Jansson and Karlberg 2011), *θ* (-):

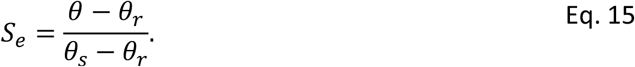

In Eq. 15, *θ*_*s*_ is the saturated soil water content and *θ*_*r*_ is the residual water content.

We calculate *k*_*rc*_ by solving Eq. 16 (Sperry and Love 2015).

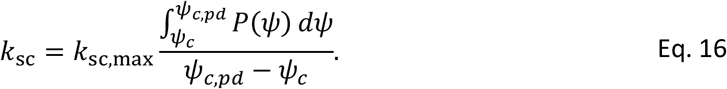

In Eq. 16, *ψ*_*c*,*pd*_ = *ψ*_*s*_ − *H ρ g* is the pre-dawn canopy water potential. We use Simpson’s 1/3 rule to approximate the integral in Eq. 16 and the equation is solved by applying a fix-point iteration method.

### Plant optimization

We define the instantaneous fitness proxy, *G* (mol m^-2^ s^-1^), as the instantaneous carbon assimilation rate at top of the canopy, 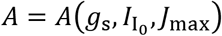, times a reduction factor *k*_cost_, representing the effect of reduced plant conductance under water stress (Eller et al. 2018), minus the cost of maintaining *J*_*max*_, i.e., (*N*_*r*_ + *N*_*u*_)*J*_*max*_. Thus,

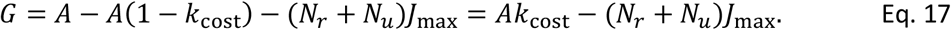

In Eq. 17, *N*_*r*_ (-) represents the leaf respiration cost as the ratio between dark respiration (*R*_*d*_) and *J*_max_, which is linked to leaf N because photosynthetic capacity and *J*_max_, increases with leaf N concentration associated with photosynthetic proteins. Because *N*_*r*_ is based on the ratio of fundamental leaf biochemical processes with similar climatic responses (Wang et al. 2020) it is relatively constant among species and climate conditions. *N*_*u*_ represents the carbon investment (fine-roots, mycorrhiza, exudation) for nutrient uptake required to construct and maintain *J*_max_ which is expected to strongly depend on soil N availability.

The costs of hydraulic risks and damage are represented by the parameter *k*_cost_ = (*k*_sc_ − *k*_crit_)/(*k*_sc,max_ − *k*_crit_) (Supplementary material of Eller et al., 2018). Based on the commonly observed lethal loss of conductivity of 88% (Liang et al. 2021) here we assumed *k*_crit_ = 0.12*k*_sc,max_ and the *A*(1 − *k*_cost_) cost term we assume that: 1) each fraction of *k*_sc_ corresponds to an equal loss in functional leaf area and thus assimilation loss and 2) in the event of hydraulic failure and fatal embolism, i.e., when *k*_*rc*_ decreases and approaches 0, the loss should be equal to the total carbon gain.

While *G* represents the instantaneous fitness reward, we assume that *N*_*m*,*f*_ and *g*_*s*_ regulate such that the accumulative fitness, i.e. the integration of *G* over time, is maximized. Furthermore, we assume that: i) *N*_*m*,*f*_ optimizes on a weekly time scale and *g*_*s*_ on a sub-daily time scale. ii) *N*_*f*_ is constant over a week’s period. iii) The day-to-day change of the weather variables within a week are neglectable compared to the within-day variation. iv) The daily integration of *G* can be approximated by the sum of two instantaneous function values according to the two-segment daily model (SDM-2, Wang et al., 2014). The original segmented daily model assumed that the nonlinear responds of *A* can be approximated by a pricewise linear function, i.e., the response curve can be approximated by a number of line segments, and that weather variables (specifically radiation, temperature, and relative humidity) follow a sine function. With these assumptions, we propose a two-step optimization routine. In the first step, we optimize *N*_*m*,*f*_ and two *g*_*s*_ values to maximize the integral of *G* over a specified week (long-term optimization). The two *g*_*s*_ values represent the within-day variation of stomatal conductance (one *g*_*s*_ value for each segment in SDM-2) for an average day within the specific week. In the second step, we maximize the daily integral of *G* for each day in the specified week by re-optimizing the two *g*_*s*_ value for each day (fine-tuning). Here we use the optimal weekly *N*_*m*,*f*_ value from the previous step as input, and the two average-day *g*_*s*_ values from the first step are used as initial guesses for the optimization algorithm. Additional information regarding the plant optimization is available in the supplementary information (*Methods S1*). In order to use the SDM-2 approximation, we need to estimate within-day values for the weather variables. To this end we use the same trigonometric functions from Wang et al. (2014) to model the diurnal change of the ambient temperature and radiation, the remaining weather variables are either calculated from these (VPD) or assumed to be constant during the day (*C*_*a*_ and *θ*), see *Methods S1* for more information.

The optimization problem is solved by using the implementation of Broyden–Fletcher–Goldfarb– Shanno algorithm (BFGS) from the *Optim*.*jl* package (K Mogensen and N Riseth 2018). We search the optimum in the range 0.007 ≤ *N*_*m*,*f*_ ≤ 0.05 and 0.001 ≤ *g*_*s*_ ≤ *g*_*s*,*crit*_ for each *g*_*s*_ and *N*_*m*,*f*_ value. Here, *g*_*s*,*crit*_ is the stomatal conductance which results in *G*(*ψ*_*r*_) = 0.12.

## Upscaling to stand-level

### Stand level primary production

We assume that *J*_max_ and *g*_s_ acclimate to irradiance resulting in a proportional relationship with irradiance level (Landsberg and Sands 2010). Then, the instantaneous gross primary production per ground area of canopy vegetation (GPP of trees), GPP_c_, is calculated as:

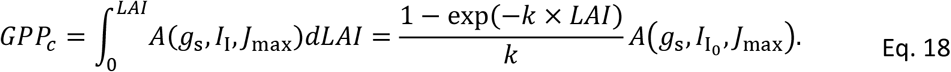

In Eq. 18, *LAI* is the leaf area index (projected area), and *k* and *I*_I_ are the light extinction coefficient and irradiance incident on a leaf, respectively, see *Methods S2* for further details. Analogue to the plant optimization routine, we employ the SDM-2 approximation (Wang et al. 2014) to calculate daily GPP value from instantaneous values. In order to compare with eddy-covariance data (ecosystem GPP, GPP_*e*_), we accounted for understory vegetation GPP, GPP_*g*_, to calculate GPP_*e*_ = GPP_*c*_ + GPP_*g*_. We assume that light use efficiency (LUE, defined as GPP/absorbed light) is the same for both vegetation layers (Tian et al. 2021). GPP_*c*_ can then be upscaled by using an upscale factor, *ζ* = GPP_*e*_/GPP_*c*_. The value of *ζ* depends on the understory vegetation LAI and corresponding light extinction coefficient (see *Methods S3* for more information). We estimate that *ζ* ≈ 1.2 for the fertilized stand and *ζ* ≈ 1.13 for the control (Table S1). These corresponding contribution of GPP_*g*_ to GPP_*e*_ was 17% and 12% for the fertilized stand and control, respectively. This is in line with previous estimates (Chi et al. 2021). Hereafter, we will refer to GPP_*e*_ as simply GPP.

### Canopy transpiration

We neglect the effects of boundary layer conductance (combined leaf and canopy boundary layer), thus the instantaneous canopy transpiration *E*_*c*_ is calculated as:

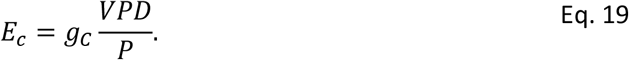

In Eq. 19, *g*_*C*_ is the canopy conductance. The canopy conductance is calculated as:

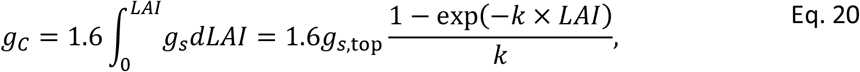

see *Methods S2* for further details. Again, we employ the SDM-2 approximation (Wang et al. 2014) to calculate the daily values for *E*_*c*_.

The data used for model calibration are based on measurements from the experimental site Rosinedal (64°10′ N, 19°45′ E). The site is a 90-year-old naturally regenerated Scots pine forest, regenerated with seed trees in 1920-1925. In 1955, the stand was pre-commercially thinned, followed by thinnings in 1976 and 1993. The experiment was established in 2005 and annual N fertilization started in 2006 with addition of 100 kg N ha^-1^ year^-1^ from 2006 to 2011, and reduced to 50 kg N ha^-1^ year^-1^ in 2012 (Lim et al. 2015). We used weather data and measured tree dimensions from the fertilized and the control stand as input for the model, see Table 1 for a full list of input variables and Fig. 2 for a depiction of the weather data time series. Briefly, stem diameter was measured at 1.3 m (DBH) annually for all trees in each of the three mensuration plots (1000m^2^) for each treatment stand. Tree height and length of live crown were measured on 20 trees per plot, using Vertex 4 Ultrasonic Hypsometer (Haglöf Inc., Sweden). We developed a relationship between height and DBH following the recommendation of Näslund (1947); parameters of the function were estimated in each year for each plot, and then applied to all individual trees. Based on destructive tree harvests in June 2006, October 2012, and October 2018, we developed allometric equations for foliage, stem and branch biomass. From 2012 and 2018 harvest samples, subsets of fresh foliage samples were scanned and dried to estimate specific leaf area. Biomass of each component was predicted based on a combination of the tree dimensions and the developed allometric equations. We estimated leaf area index by multiplying foliage biomass and specific leaf area estimates. Model outputs were calculated using daily data and validated against the eddy-covariance based ecosystem *GPP* estimates (Zhao et al. 2022) and *E*_*C*_ values for the growth periods of 2015-2018. The growing seasons was assumed to start when daily mean temperature ≥ 5 °C for five successive days, and end when daily mean temperature was < 5 °C for five successive days. The *E*_*C*_ values are estimated using the empirical model from Tor-ngern et al. (2017) with corresponding parameter estimates for the study site, provided in that paper.

**Fig. 2.**
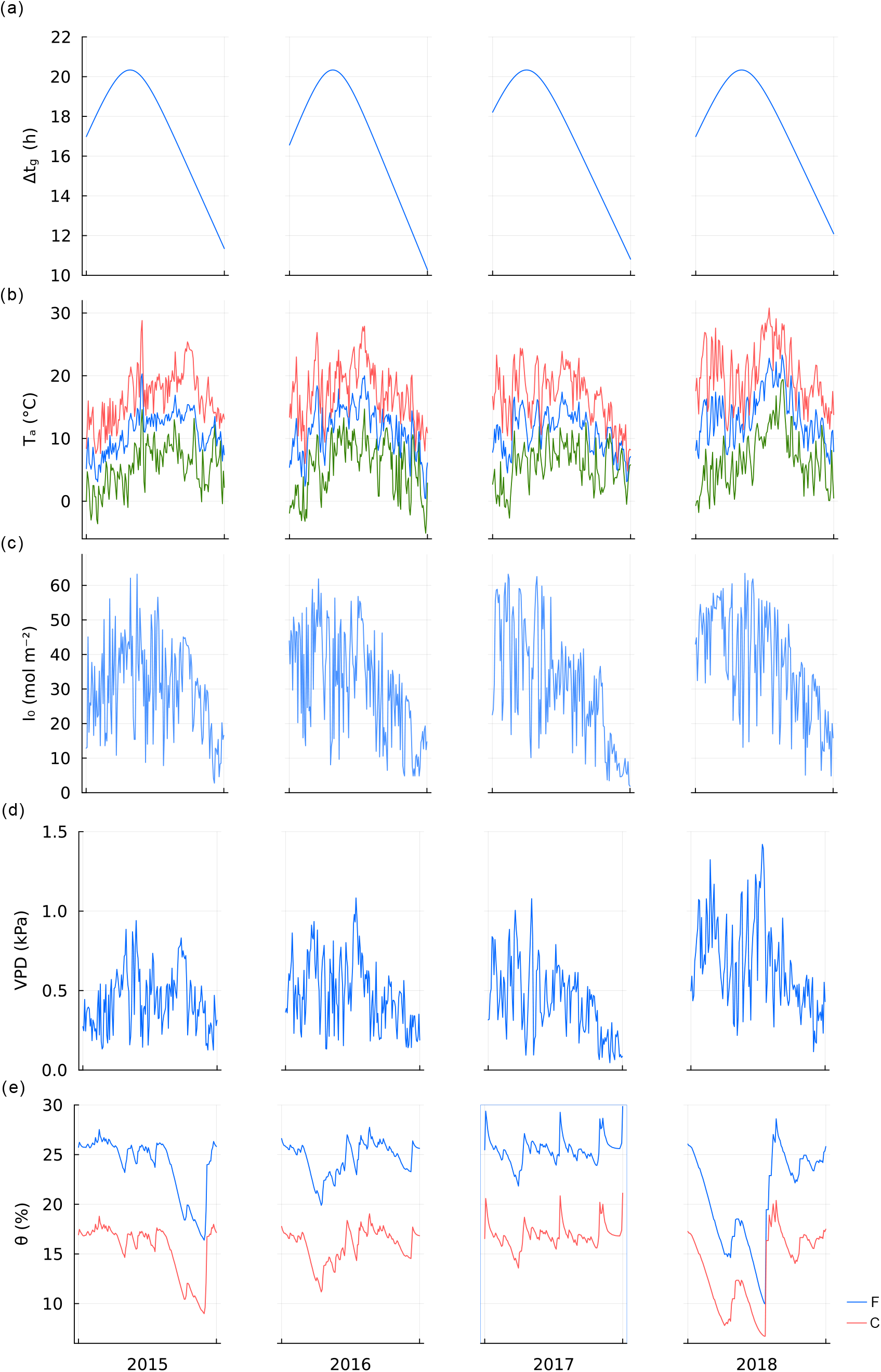
Weather data for the fertilized (F) and control (C) stand at the experimental site Rosinedal during the growth period between 2015-2018. Climate drivers: daylight hours (Δt_g_) (a), ambient air temperature (T_a_) (daily max, daily min, and daily mean) (b), daily photosynthetic active radiance (I_0_) (c), vapor pressure deficit (VPV) (d), soil volumetric water content (θ) of the fertilized stand (C) and the control (C). These values were used as input for our model.

### Parameter estimation

We estimated the unknown set of parameters of the model, ***θ*** (see Table 3 for a full list of estimated parameters), by fitting the model to observations. The remaining model parameters were taken from other sources, see Table 2. To minimize the effect of data outliers, we assume that model and measurement errors are Laplace distributed (Tian et al. 2021), i.e., *y*_*i*,*j*,*k*_ − *M*(*X*_*j*,*k*_, ***θ***)_*i*,*j*,*k*_ = *e*_*i*,*j*,*k*_∼*Laplaceo*, (0, *a*_*i*_ + *b*_*i*_*M*(***θ***)_*i*,*j*,*k*_). Here, *y*_*i*,*j*,*k*_ denotes the response variables (the measured variables), *X*_*j*,*k*_ are the collection of explanatory variables (tree size and climate data). *M* is the model output, *e*_*i*,*j*,*k*_ are the model errors, *Laplace*(*μ, b*) is the Laplace distribution with location parameter *μ* and scale parameter *b* > 0. The parameters *a*_*i*_ and *b*_*i*_ control the variance of the error distribution and we assume that the scale parameter is linear with respect to the model output (Tian et al. 2021). We also use data type specific weights, *w*_*i*_, to weigh the importance of different data types. The subscripts *i, j, k* denotes the data type (*GPP, E*_*C*_, or *N*_*m*,*f*_), the stand treatment (control or fertilized), and the data index (individual observations), respectively. This assumption leads to the following likelihood function,

**Table 2:** Parameter values used in the model.

| Parameter | Description | Value (units) | Reference/Note |
| --- | --- | --- | --- |
| *Gross primary production model parameters* |  |  |  |
| $\Gamma_{ref}^*/E_{A,\Gamma}$ | The CO<sub>2</sub> compensation point at 25 °C/Activation energy of $\Gamma^*$ | 4.17 (Pa)/ 23.42 (kJ mol<sup>-1</sup>) | Table 3.1 in Landsberg & Sands (2010) |
| $E_{A,J}/E_{D,J}/T_{opt}$ | Activation energy of $J_{max}$/ Deactivation energy of $J_{max}$/Optimal temperature for $J_{max}$ | 47.4 (kJ mol<sup>-1</sup>) / 200 (kJ mol<sup>-1</sup>) / 305 (K) | (Tarvainen et al. 2018) |
| $\theta_j$ | Curvature of the light response curve | 0.7 (-) | Table 3.1 in Landsberg & Sands (2010) |
| $k$ | light extinction coefficient | 0.52 (-) | (Tian et al. 2021) |
| $m$ | Leaf transmittance | 0.05 (-) | ch. 5.1.1 in Landsberg & Sands (2010) |
| *Hydraulics model parameters* |  |  |  |
| $\psi_{50,sc}/b_{sc}$ | water potential which causes 50% loss in soil-canopy hydraulic conductivity/Sensitivity of soil-canopy hydraulic conductivity to water potential | -2.7 (MPa) / 2.15 (-) | $\psi_{50,sc}$ and $b_{sc}$ are estimated from Aguade et al. (2015) |
| $\theta_s/\theta_r$ | Saturated soil water content/ Residual water content | 0.41 (-) / 0.006 (-) | From unpublished Pf-curve |
| $\psi_a/\lambda$ | air-entry tension/pore size distribution index | -0.098 (MPa) / 1 (-) | From unpublished Pf-curve |
| *Nitrogen cost parameter* |  |  |  |
| $N_r$ | Ratio between dark respiration and $J_{max}$ | 0.0056 (-) | Estimated from Kellomäki and Wang (1997) |
| *Variance parameters* |  |  |  |
| $a_{E_c,j}/b_{E_c,j}$ | The parameters which control the variance of the error distribution for $E_c$ (mm day<sup>-1</sup>/-) | 0.069/0.15 ($j$ = Fertilized) 0.043/0.18 ($j$ = Control) | Tian et al., 2021 |
| $y_{N_{m,f,j,k}}/a_{N_{m,f,j}}/b_{N_{m,f,j}}$ | The response variable and parameters which control the variance of the error distribution for $N_{m,f}$ (Kg Kg<sup>-1</sup>/ Kg Kg<sup>-1</sup>/-) | 0.021/0.0023/0 ($j$ = Fertilized and for all $k$) 0.012/0.0023/0 ($j$ = Control and for all $k$) | Estimated from data (Lim et al. 2015) |

**Table 3:**
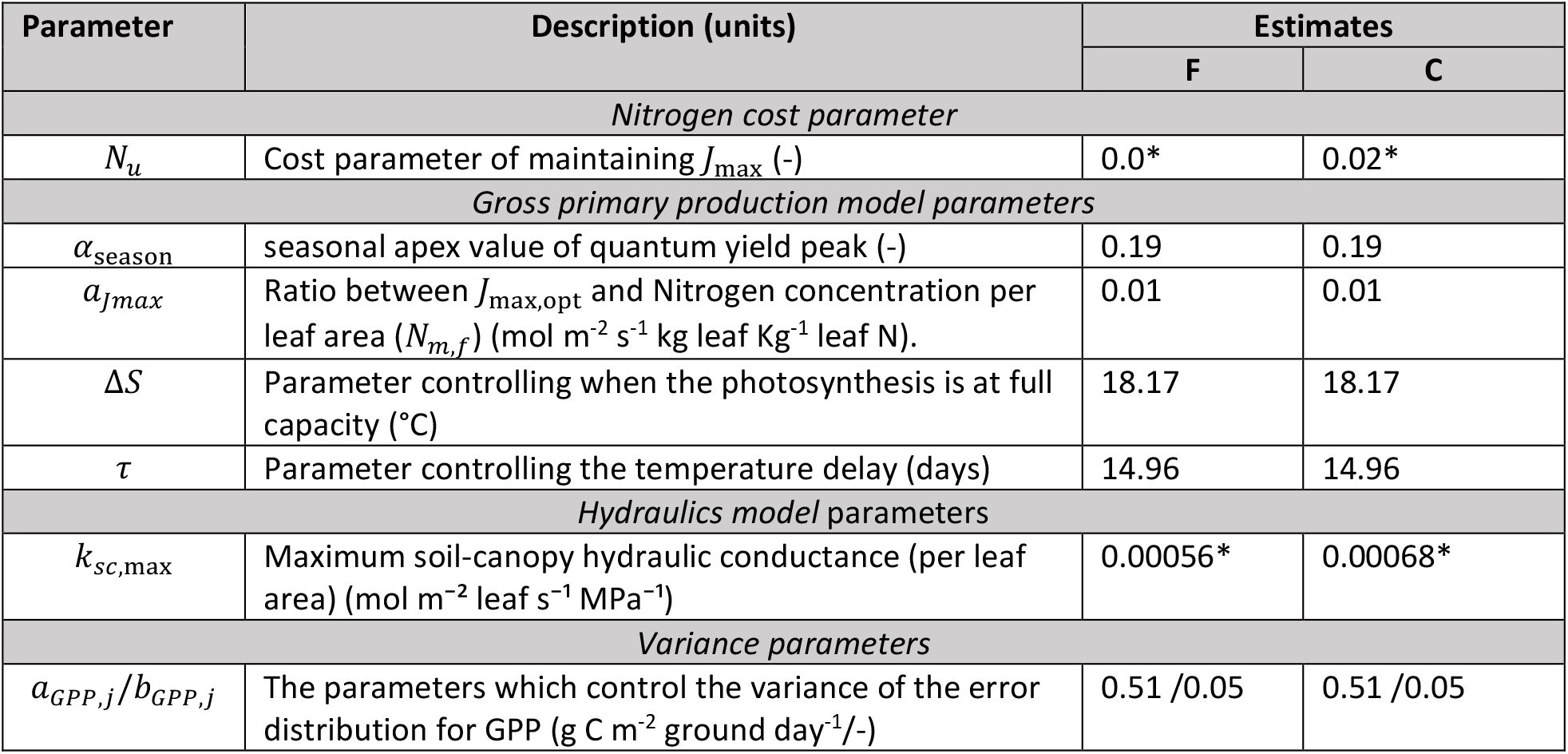
Estimated parameters. All parameters were shared among the stand treatments (fertilized F and control C), except when indicated by an asterisk (*).

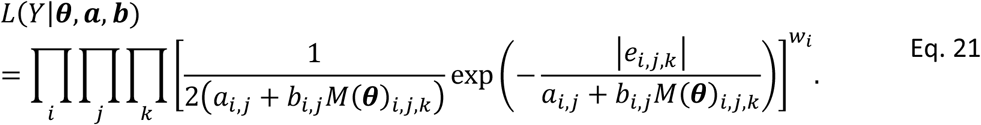

In Eq. 21, ***a, b*** are collections of *a*_*i*_ and *b*_*i*_ values, respectively. The values for 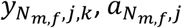, and 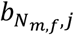, were estimated from data Lim et al. (2015) and the values of 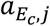 and 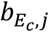 were taken from Tian et al. (2021) (Table 2), while *a*_*GPP*,*j*_ and *b*_*GPP*,*j*_ were estimated in conjunction with the unknown model parameters and we used 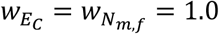 and *w*_GPP_ = 1.5.

The parameter estimates, 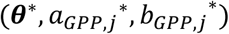, were determined by maximization of the likelihood function, i.e., 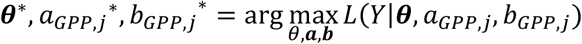. To find the maximum, we employed the adaptive differential evolution optimizer, a global optimization algorithm, from the *BlackBoxOptim*.*jl* package (Feldt 2018).

We performed two parameter estimation and validation cases: One where all the parameters in Table 3 are shared between the two stand treatments, i.e., same parameter values were used for both treatments, with the exception of *N*_*s*_ and *k*_*rc*,max_. In the second case, none of the parameters in Table 3 were shared between the two stand treatments, i.e., the parameters were estimated separately for the control and fertilized stand. The result of the first case will be shown in the *Results* section and the result of the second case can be viewed in *Methods S4* and Fig. S3-Fig. S5.

### Model validation

To validate our model, we excluded 20% of the datapoints (119 out of 588 datapoints) to form a validation dataset. The validation dataset was generated by randomly selecting datapoints from the complete dataset. The remaining datapoints, the training dataset, were used for parameter estimation. To minimize the effect of selection bias, we repeated the parameter estimation and validation process 10 times for both cases, thus creating 10 random validation and training datasets.

## Results

### The model can predict seasonal changes in *GPP* and *E*_*C*_

The result for the parameter estimation is depicted in Fig. 3. Estimated parameters are provided in Table 3. The R^2^ for all datapoints (training set + validation set), for the run with the highest likelihood, was 0.73 (*GPP*) and 0.82 (*E*_*C*_) for the fertilized stand. The corresponding values for the control were 0.71 (*GPP*) and 0.81 (*E*_*C*_) (Table 4). (error estimates for all 10 runs are depicted in Fig. S2). Overall, our model was able to capture the inter-seasonal variation of *GPP* and *E*_*c*_.

**Table 4:**
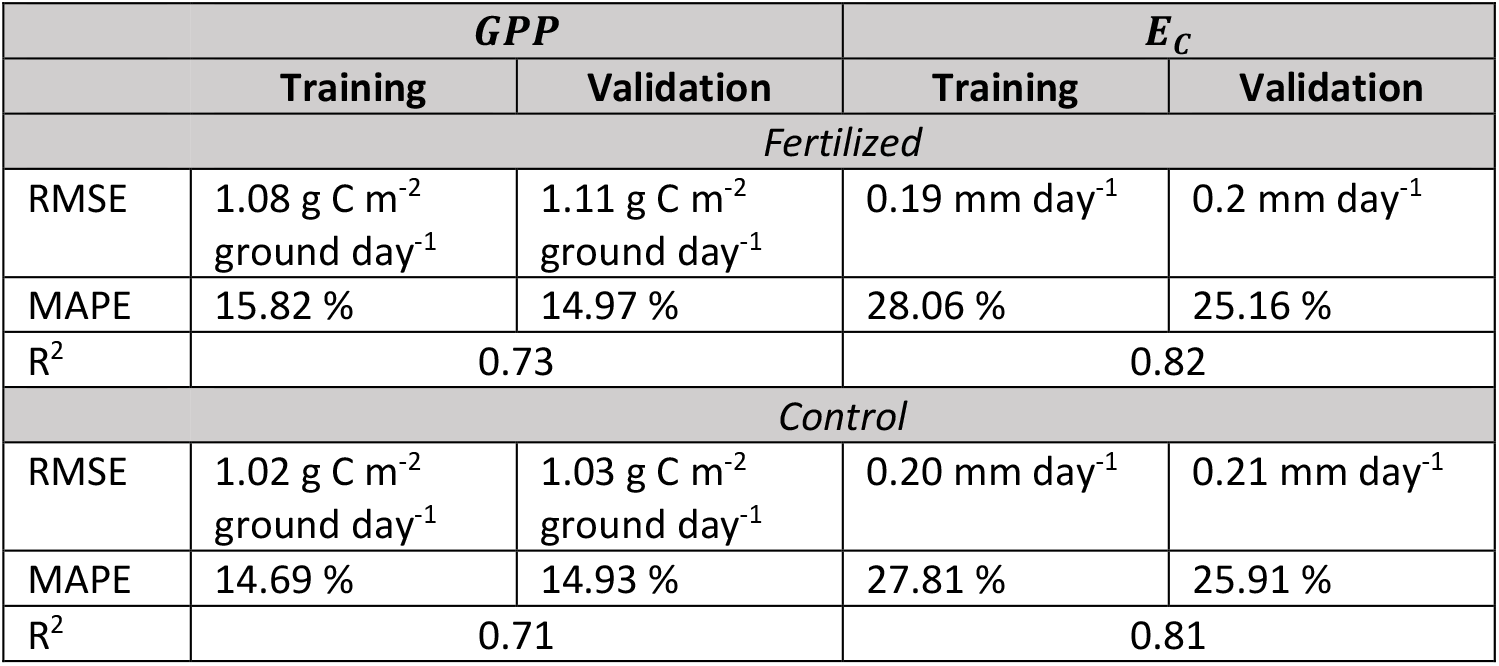
Model Performance measure (root-mean-square error, RMSE; Mean absolute percentage error, MAPE; Coefficient of determination R2) during the parameter estimation (training) and model validation (validation). Parameters where shared among both stand treatments (fertilized and control). Note, R2 was calculated over all datapoints (training + validation).

**Fig. 3.**
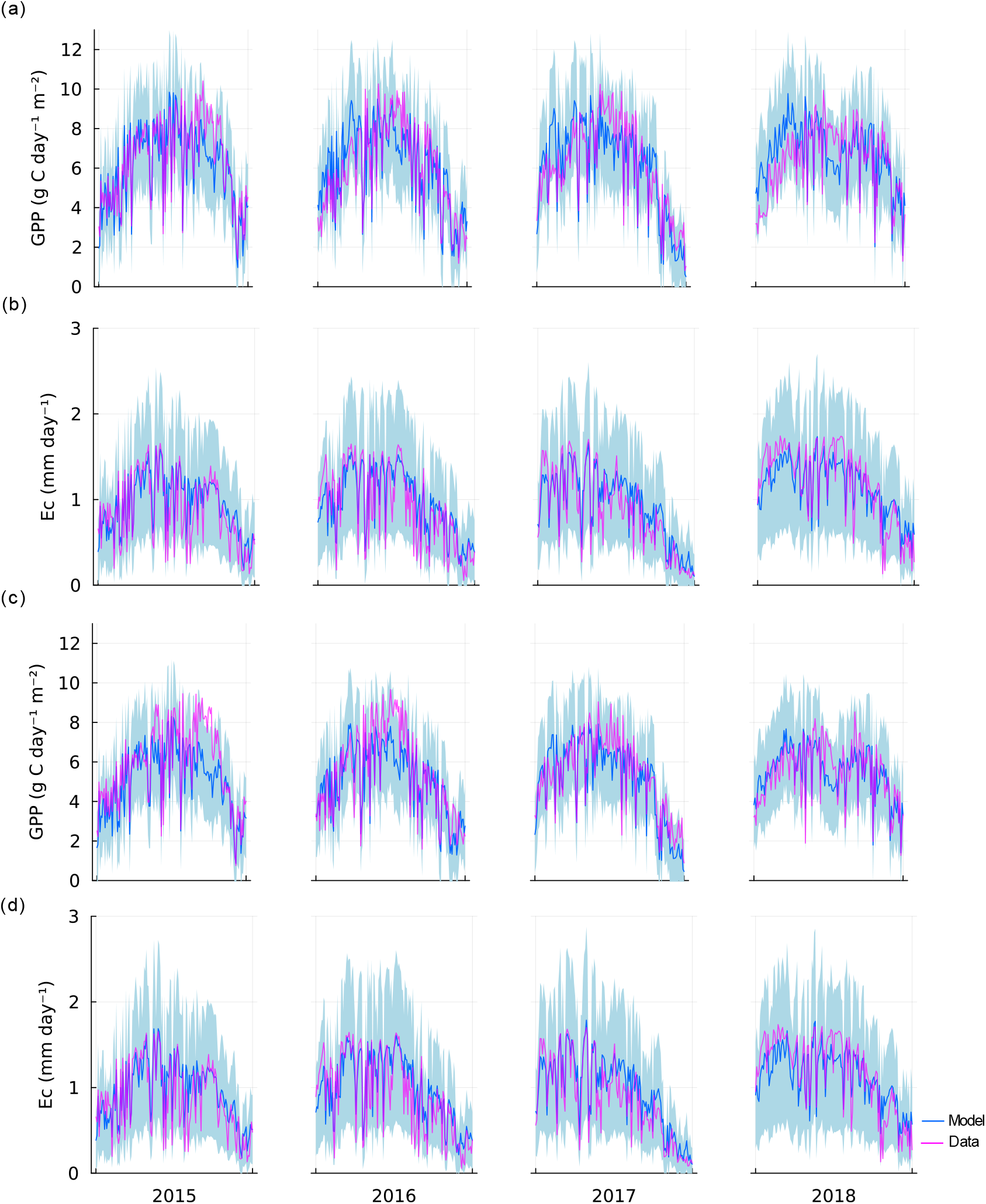
Result from the parameter estimation case when most of the parameters are shared between the two stands, fertilized (a-b) and control (c-d). (a) and (c), and (b) and (d) depicts the data against the corresponding simulated values for ecosystem gross primary production (GPP) and canopy transpiration (E_C_), respectively. The 95% confidence interval of the fitted Laplace distribution (see Parameter estimation section) is depicted by the shaded area.

### The effect of soil N availability on *GPP* and *E*_*C*_ is captured by hydraulic conductance per leaf area and the site-specific nitrogen cost

The summary statistics of the shared and non-shared parameter estimation case showed similar predictive performance. For the non-shared parameter case, the R^2^ was 0.73 (*GPP*) and 0.82 (*E*_*C*_) for the fertilized plot and 0.75 (*GPP*) and 0.81 (*E*_*C*_) for the control plot (see *Methods* S*4* and Table S2). The result suggests that the difference between the two treatments can be well captured by the differences in N acquisition cost (*N*_*u*_) and hydraulic conductivity per leaf area (*k*_sc,max_). *N*_*u*_ (unitless) was 0.00 and 0.02 and *k*_sc,max_ was 0.56 and 0.68 mmol m^-2^ leaf s^-1^ MPa^-1^ for the fertilized and control treatments, respectively (Table 3).

### Environmental drivers of leaf N concentration, stomatal conductance, and water use efficiency

Fig. 4 illustrates the variation in optimal stomatal conductance (*g*_*s*_) and leaf N concentration (*N*_*m*,*f*_) with respect to the weather variables including irradiance (*I*_0_), mean ambient temperature (*T*_*a*_), vapor pressure deficit (VPD), and soil water content (*θ*). We calculated the Pearson correlation coefficients between the plant variables and the weather variables. For *N*_*m*,*f*_ the correlation was 0.01 (*P*=.77), -0.70 (*P*<.001), -0.26 (*P*<.001), and 0.30 (*P*<.001) for *I*_0_, *T*_*a*_, VPD, and *θ*, respectively. For *g*_*s*_ the corresponding values were -0.56 (*P*<.001), -0.43 (*P*<.001), -0.77 (*P*<.001), 0.32 (*P*<.001). The result indicates that the optimal value of *N*_*m*,*f*_ mostly respond to changes in *T*_*a*_, whereas *g*_*s*_, responds mostly to the change in VPD and *I*_0_. Correlation values shown here are for the fertilized stand. The control showed similar results and these values can be viewed in Table S3.

**Fig. 4.**
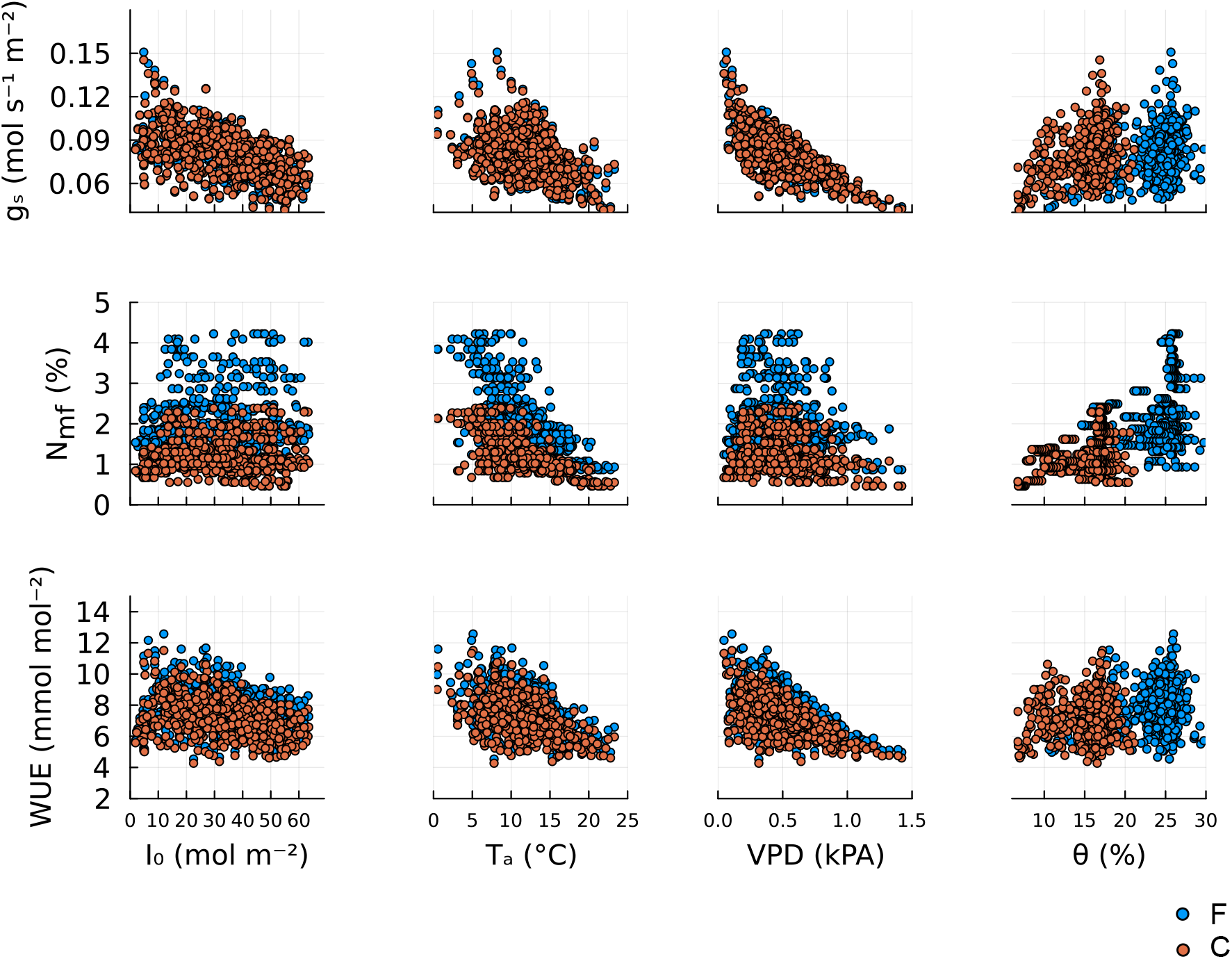
Response of the optimal, stomatal conductance (g_s_, first row), leaf mass-based N concentration (N_m,f_, second row), and water use efficiency (WUE) to change in weather variables: above canopy photosynthetic active radiance (I_0_, first column), mean ambient air temperature (T_a_, second column), vapor pressure deficit (VPV, third column), and soil water content (θ, fourth column). The blue circles correspond to the optimal trait values for the fertilized stand (F) and orange circles correspond to the control stand (C). Correlation values between traits and weather variables can be found in Table S3. The figure was generated by applying the model to the environmental data (Fig. 2), using parameters from Table 2 and Table 3.

The response of water use efficiency, wue = GPP/*E*_*c*_, to meteorological variables *I*_0_,*T*_*a*_, VPD, and *θ* are shown in Fig. 4; here, GPP reference to the gross primary production of canopy vegetation. The correlation coefficients between wue and meteorological variables were -0.29 (*P*<.001), -0.47 (P<.001), -0.57 (*P*<.001), 0.15 (*P*<.001) for *I*_0_, *T*_*a*_, VPD, and *θ*, respectively. From the correlation values and Fig. 4 we see that wue responds similarly to the various weather variables as *g*_*s*_. In contrast, the wue response showed little resemblance to the weather response of *N*_*m*,*f*_, indicating that wue is mainly influenced by *g*_*s*_.

## Discussion and conclusions

### Model scope and limitations

Our model estimates canopy transpiration (*E*_*c*_) and GPP, based on optimal acclimation of stomatal conductance and leaf nitrogen (N) concentration. The model can, for the most part, accurately predict observed inter- and intra-seasonal variation of *E*_*c*_ and GPP at the two study sites (Fig. 3). An exception is the growth period of 2018, where the site was hit by a severe drought period. During this period the model overestimates GPP for both the fertilized stand and the control. This divergence may be linked to the fact that we use a fixed potential hydraulic conductivity, i.e., not accounting for hydraulic damages, see below. Our results demonstrate that the effect of fertilization can be captured by adjustments in only two parameters: maximum soil-to-canopy hydraulic conductance per leaf area, *k*_sc,max_, and the site-specific N acquisition cost parameter, *N*_*s*_. The current version of the model predicts GPP and transpiration for trees with a given LAI and potential hydraulic conductivity per LAI (*k*_sc,max_). As such, it is naturally unable to predict dynamics of these properties, which could be addressed in future versions of the model (see Conclusions and Outlook, below).

### The impact of increased soil N availability

The difference between our model and the previous models by Eller et al. (2018) and Prentice et al. (2014) is that we do not only account for optimal stomatal response but also optimize leaf N concentration. Another recent model that allows optimization of stomatal conductance and *J*_max_, which is functionally equivalent to our optimization of leaf N, is the model by Joshi et al. (2022). However, this model has a different hydraulic cost function and, more importantly, does not address variation in the cost of N uptake related to soil N availability.

Our model allows us to account for the response to increased soil N availability. In agreement with our model predictions, it has been observed that increased soil N availability results in increased leaf N concentration (Lim et al. 2015, Tarvainen et al. 2016). This, in turn, has a positive impact on the potential photosynthetic capacity and the *J*_max_ in our model. The model results suggest that the fertilization treatment radically reduced the trees’ C cost for N uptake (the unitless cost parameter *N*_*u*_) from 0.017 to 0.0. However, without further empirical evidence, the zero cost of N uptake should be interpreted with care, rather as being too low to be separated from other costs by the model analysis than as an absolute zero value. Nevertheless, the large difference in N uptake costs agrees with the concurrent observed difference in C allocation to the components contributing to N uptake: fine-root, mycorrhiza, and exudates production (Marshall et al. 2023). The low-cost uptake of N may be possible via mass flow of dissolved N in water uptake, which can be the dominant process of N uptake in fertilized conditions (Oyewole et al. 2017, McMurtrie and Näsholm 2018, Henriksson et al. 2021) where the chemical profile of soil N is biased towards mineral forms (e.g. nitrate) that are mobile in the soil solution. This effect may have been further enhanced by higher soil water in the fertilized plot related a higher field capacity and more organic matter than in the control plot (Tian et al. 2021). Other than fertilized soils, this may occur under specific conditions where soil nitrification rates are high. In contrast, the control stand represents a more common boreal forest soil profile, where N is present mainly in organic and less mobile forms which drive up the N acquisition cost.

Besides the decrease in *N*_*u*_, we also get an increase in LAI in the fertilized plot. Because the sapwood cross sectional area has not increased to the same extent, we get a 12% lower Huber value (sapwood area / leaf area) in the fertilized compared with the control stand (calculated from field measurements). This contributes to the estimated 17% decrease in estimated conductivity per leaf area (*k*_*sc*,max_) in the fertilized plot. These results may reflect an optimality response to a lower cost of N uptake and thus lower cost of leaf area.

### The responses of leaf variables and water use efficiency to weather variables

Based on our results, we can infer that the optimal leaf N concentration (*N*_*m*,*f*_) is negatively correlated with temperature (Fig. 4). This is in line with empirical observations for Scots pine (Zha et al. 2002), understory evergreen plants (Muller et al. 2011), and a global biogeographic pattern of different plant species (Reich and Oleksyn 2004). This is caused by the shape of the C assimilation curve and the cost of maintaining *J*_max_ as a function of *N*_*m*,*f*_. The optimum *N*_*m*,*f*_ will occur when the slope of the C assimilation function is equal to the slope of the cost line. When the temperature increases, the slope of the cost line increases more than the C assimilation curve, and thus the optimal *N*_*m*,*f*_ will decrease (Fig. S8). This temperature effect further implies that the leaf N content should decrease with higher temperatures in the middle of the growth season (see Fig. S7). Because the model only accounts for photosynthetic N but no other forms (e.g., N for structural purposes or storage) that may respond differently, it probably overestimates the seasonal N variation. Furthermore, the model does not consider seasonal dynamics in the vertical N distribution in the canopy. Nevertheless, qualitatively similar seasonal variation has been observed in a nearby boreal Scots pine forest (Näsholm and Ericsson 1990) as well as a temperate Scots pine forest (Wyka et al. 2016).

As expected, we found that stomatal *g*_*s*_ correlated best with VPD. A high VPD leads to an increase in water loss and negative water potential for a fixed value of *g*_*s*_. Thus, *g*_*s*_ is reduced in response to rising VPD, in order to reduce the hydraulic risk cost. Our results (Fig. 4) further show that the stomatal *g*_*s*_ is negatively correlated to the above canopy photosynthetic active radiance (*I*_0_). However, this correlation does not correspond to a direct relationship but reflects a correlation between VPD and *I*_0_ and confounding variation in other drivers, which was confirmed by running the model with fixed weather variables (except for *I*_0_). Furthermore, we found that the response of wue to the environmental variables closely follows that of *g*_*s*_ (Fig. 4). Thus, wue is more strongly correlated to *g*_*s*_ than *N*_*m*,*f*_, which is not surprising as water used as transpiration is directly regulated by *g*_*s*_. In all cases, soil water content (*θ*) had a small impact on *g*_*s*_, *N*_*m*,*f*_, and wue when *θ* is large (approximately *θ* > 25% for F treatment and *θ* > 15% for C treatment). For lower soil water content, a decrease in *θ* leads to decreases in *g*_*s*,top_, *N*_*m*,*f*,top_, and wue, driving the observed positive correlation between *θ* and the plant variables and process rates. The difference between high and low *θ* is caused by the monotonically decreasing concave down shape of the vulnerability function, *G*(*ψ*). For low values of *θ* (highly negative soil water potential, *ψ*_*s*_), the increase in cost associated with small changes in plant variables (the derivative of the cost with respect to the plant variables) is larger than for higher *θ*.

In summary, while both *g*_*s*_ and *N*_*m*,*f*_ correlated well with different weather variables there is no significant correlation between the two plant variables (Pearson correlation = -0.017, *p*=0.68 for F treatment and -0.0026, *p*= 0.95 for C treatment, Fig. S6). This implies that *g*_*s*_ and *N*_*m*,*f*_ mostly respond to different meteorological variables. *g*_*s*_ responds strongly to irradiance and VPD, while *N*_*m*,*f*_ responds strongly to the ambient temperature.

### Outlook

The key scientific advancement made by this improved model lies in its ability to explain, and accurately predict the interacting effects of climate, soil water availability, and soil N availability on GPP and transpiration, based on an eco-evolutionary optimality principle (EEO). This capacity makes the model well suited for studying the impact of climate change in N limited boreal forests. Also, the effects of changes in soil N availability such as fertilization treatments and N deposition can be studied by our model. However, one has to be careful when applying the model to severe drought conditions as it does not yet account for accumulation of hydraulic damages. To address long-term effects on forest growth, the model can be extended with the capacity to predict carbon allocation to the different plant organs, such as branches, fine root, stem, and leaves. To this end, our model could be coupled with allocation models based on similar EEO principles (e.g., Franklin et al., 2012 and Fransson et al., 2021).

## Acknowledgement

The study site Rosinedal is part of the Swedish Infrastructure for Ecosystem Science (SITES) and financial support from the Swedish Research Council (VR) and contributing research institutes to SITES are acknowledged.

## Author contributions

O.F. and P.F conceived the study and formulated the model. P.F. implemented and analyzed the model, and wrote the first draft of the manuscript. H.L. provided biometric data, P.Z. and M.P. provided the eddy covariance data. All authors discussed the results and implications and revised the manuscript.

## Supplementary Data

**Supplementary Data**

**Methods S1**. Details regarding the plant optimization routine.

**Methods S2**. Describing how to upscale *GPP* and *E*_*C*_ values from leaf-level to stand-level.

**Methods S3**. Description of how to estimate the contribution of understory vegetation to ecosystem *GPP*.

**Methods S4**. Result from the parameter estimation case when no shared parameters between the stands.

**Fig. S1**. Statistical summaries of the result from parameter estimation for the parameter estimation case when parameters are shared among for the fertilized stand and control stand.

**Fig. S2**. Model estimates of *GPP* and *E*_*C*_, from the parameter estimation case when no parameters are shared between the fertilized stand and control, against data.

**Fig. S3**. Statistical summaries of the result from parameter estimation for the parameter estimation case when no parameters are shared among for the fertilized stand and control stand.

**Fig. S4**. Optimal stomatal conductance (*g*_*s*_) against optimal leaf mass-based N concentration (*N*_*m*,*f*_).

**Fig. S5**. Optimal stomatal conductance (*g*_*s*_) and optimal leaf mass-based N concentration (*N*_*m*,*f*_). against time.

**Fig. S6**. Carbon assimilation curve against cost line of maintaining *J*_max_.

**Table S1**. Canopy *GPP* scaling factor and the fraction contribution of ground vegetation layer to eco system *GPP*.

**Table S2**. Estimated parameter values for the case when no parameters are shared between the two stand treatments.

**Table S3**. Pearson correlation between model outputs and weather variables.

## References

Aguade D, Poyatos R, Gomez M, Oliva J, Martinez-Vilalta J (2015) The role of defoliation and root rot pathogen infection in driving the mode of drought-related physiological decline in Scots pine (Pinus sylvestris L.). Tree Physiol 35:229–242. https://academic.oup.com/treephys/article-lookup/doi/10.1093/treephys/tpv005

Alduchov OA, Eskridge RE (1996) Improved Magnus Form Approximation of Saturation Vapor Pressure. Journal of Applied Meteorology 35:601–609.

Anderegg WRL, Wolf A, Arango-Velez A, Choat B, Chmura DJ, Jansen S, Kolb T, Li S, Meinzer FC, Pita P, Resco de Dios V, Sperry JS, Wolfe BT, Pacala S (2018) Woody plants optimise stomatal behaviour relative to hydraulic risk. Ecol Lett 21:968–977.

Ball JT, Woodrow IE, Berry JA (1987) A Model Predicting Stomatal Conductance and its Contribution to the Control of Photosynthesis under Different Environmental Conditions. In: Progress in Photosynthesis Research.

Binkley D, Högberg P (2016) Tamm Review: Revisiting the influence of nitrogen deposition on Swedish forests. For Ecol Manage 368:222–239.

Chen JL, Reynolds JF, Harley PC, Tenhunen JD (1993) Coordination theory of leaf nitrogen distribution in a canopy. Oecologia 93:63–69.

Chi J, Zhao P, Klosterhalfen A, Jocher G, Kljun N, Nilsson MB, Peichl M (2021) Forest floor fluxes drive differences in the carbon balance of contrasting boreal forest stands. Agric For Meteorol 306

Cowan IR, Farquhar GD (1977) Stomatal function in relation to leaf metabolism and environment. Symp Soc Exp Biol 31

Eller CB, Rowland L, Oliveira RS, Bittencourt PRL, Barros F V., Da Costa ACL, Meir P, Friend AD, Mencuccini M, Sitch S, Cox P (2018) Modelling tropical forest responses to drought and El Niño with a stomatal optimization model based on xylem hydraulics. Philosophical Transactions of the Royal Society B: Biological Sciences 373

Farquhar GD, von Caemmerer S (1982) Modelling of Photosynthetic Response to Environmental Conditions. In: Physiological Plant Ecology II.

Farquhar GD, von Caemmerer S, Berry JA (1980) A biochemical model of photosynthetic CO2 assimilation in leaves of C3 species. Planta 149:78–90. https://link.springer.com/article/10.1007/BF00386231%0Apapers3://publication/doi/10.1007/BF00386231

Feldt R (2018) BlackBoxOptim.jl. GitHub repository

Flo V, Joshi J, Sabot M, Sandoval D, Prentice IC (2023) Incorporating photosynthetic acclimation improves stomatal optimisation models. bioRxiv

Franklin O (2007) Optimal nitrogen allocation controls tree responses to elevated CO 2. New Phytologist 174:811–822.

Franklin O, Johansson J, Dewar RC, Dieckmann U, McMurtrie RE, Brännström Å, Dybzinski R (2012) Modeling carbon allocation in trees: A search for principles. Tree Physiol 32:648–666.

Fransson P, Brännström Å, Franklin O (2021) A tree’s quest for light-optimal height and diameter growth under a shading canopy. Tree Physiol 41:1–11.

Friend AD (1991) Use of a model of photosynthesis and leaf microenvironment to predict optimal stomatal conductance and leaf nitrogen partitioning. Plant Cell Environ 14

Hari P, Mäkelä A (2003) Annual pattern of photosynthesis in Scots pine in the boreal zone. Tree Physiol 23:145–155.

Henriksson N, Lim H, Marshall J, Franklin O, McMurtrie RE, Lutter R, Magh R, Lundmark T, Näsholm T (2021) Tree water uptake enhances nitrogen acquisition in a fertilized boreal forest – but not under nitrogen-poor conditions. New Phytologist 232

Högberg P, Näsholm T, Franklin O, Högberg MN (2017) Tamm Review: On the nature of the nitrogen limitation to plant growth in Fennoscandian boreal forests. For Ecol Manage 403

IPCC (2014) IPCC Fifth Assessment Synthesis Report-Climate Change 2014 Synthesis Report. IPCC Fifth Assessment Synthesis Report-Climate Change 2014 Synthesis Report

Jansson P-E, Karlberg L (2011) Coupled Heat and Mass Transfer Model for Soil-Plant-Atmosphere Systems. Stockholm, Sweden.

Jenkins A (2013) The Sun’s position in the sky. Eur J Phys 34:633–652.

Joshi J, Stocker BD, Hofhansl F, Zhou S, Dieckmann U, Prentice IC (2022) Towards a unified theory of plant photosynthesis and hydraulics. Nat Plants 8:1304–1316.

K Mogensen P, N Riseth A (2018) Optim: A mathematical optimization package for Julia. J Open Source Softw 3:615.

Kellomäki S, Wang KY (1997) Photosynthetic responses of Scots pine to elevated CO2 and nitrogen supply: Results of a branch-in-bag experiment. Tree Physiol 17

Landsberg JJ, Sands P (2010) Physiological ecology of forest production: principles, processes and models. Academic Press.

Liang X, Ye Q, Liu H, Brodribb TJ (2021) Wood density predicts mortality threshold for diverse trees. New Phytologist 229

Lim H, Oren R, Palmroth S, Tor-ngern P, Mörling T, Näsholm T, Lundmark T, Helmisaari HS, Leppälammi-Kujansuu J, Linder S (2015) Inter-annual variability of precipitation constrains the production response of boreal Pinus sylvestris to nitrogen fertilization. For Ecol Manage 348:31–45. 10.1016/j.foreco.2015.03.029

Maire V, Martre P, Kattge J, Gastal F, Esser G, Fontaine S, Soussana JF (2012) The coordination of leaf photosynthesis links C and N fluxes in C3 plant species. PLoS One 7

Mäkelä A, Hari P, Berninger F, Hänninen H, Nikinmaa E (2004) Acclimation of photosynthetic capacity in Scots pine to the annual cycle of temperature. Tree Physiol 24:369–376.

Mäkelä A, Pulkkinen M, Kolari P, Lagergren F, Berbigier P, Lindroth A, Loustau D, Nikinmaa E, Vesala T, Hari P (2008) Developing an empirical model of stand GPP with the LUE approach: Analysis of eddy covariance data at five contrasting conifer sites in Europe. Glob Chang Biol 14:92–108.

Marshall JD, Tarvainen L, Zhao P, Lim H, Wallin G, Näsholm T, Lundmark T, Linder S, Peichl M (2023) Components explain, but do eddy fluxes constrain? Carbon budget of a nitrogen-fertilized boreal Scots pine forest. New Phytologist 239

McDowell N, Pockman WT, Allen CD, Breshears DD, Cobb N, Kolb T, Plaut J, Sperry J, West A, Williams DG, Yepez EA (2008) Mechanisms of plant survival and mortality during drought: Why do some plants survive while others succumb to drought? New Phytologist 178

McMurtrie RE, Näsholm T (2018) Quantifying the contribution of mass flow to nitrogen acquisition by an individual plant root. New Phytologist 218:119–130.

Muller O, Hirose T, Werger MJA, Hikosaka K (2011) Optimal use of leaf nitrogen explains seasonal changes in leaf nitrogen content of an understorey evergreen shrub. Ann Bot 108:529–536. https://academic.oup.com/aob/article-lookup/doi/10.1093/aob/mcr167

Näsholm T, Ericsson A (1990) Seasonal changes in amino acids, protein and total nitrogen in needles of fertilized Scots pine trees. Tree Physiol 6:267–281. https://academic.oup.com/treephys/article-lookup/doi/10.1093/treephys/6.3.267

Näslund M (1947) Empirical formulae and tables for determining the volume of standing trees: Scots pine, Norway spruce and birch in southern Sweden and in the whole of the country. Meddelanden Fran Statens Skogsforskningsinstitut, 36, 81 (in Swedish)

Oyewole OA, Näsholm T, Jämtgård S, Näsholm T, Inselsbacher E (2017) Incorporating mass flow strongly promotes N flux rates in boreal forest soils. Soil Biol Biochem 114

Prentice IC, Dong N, Gleason SM, Maire V, Wright IJ (2014) Balancing the costs of carbon gain and water transport: Testing a new theoretical framework for plant functional ecology. Ecol Lett 17:82–91.

Reich PB, Oleksyn J (2004) Global patterns of plant leaf N and P in relation to temperature and latitude. https://www.worldclimate.com

Ryan MG (2011) Tree responses to drought. Tree Physiol 31:237–239. https://academic.oup.com/treephys/article-lookup/doi/10.1093/treephys/tpr022

Sabot MEB, De Kauwe MG, Pitman AJ, Ellsworth DS, Medlyn BE, Caldararu S, Zaehle S, Crous KY, Gimeno TE, Wujeska-Klause A, Mu M, Yang J (2022) Predicting resilience through the lens of competing adjustments to vegetation function. Plant Cell Environ 45:2744–2761. https://onlinelibrary.wiley.com/doi/10.1111/pce.14376

Sabot MEB, De Kauwe MG, Pitman AJ, Medlyn BE, Ellsworth DS, Martin-StPaul NK, Wu J, Choat B, Limousin JM, Mitchell PJ, Rogers A, Serbin SP (2022) One Stomatal Model to Rule Them All? Toward Improved Representation of Carbon and Water Exchange in Global Models. J Adv Model Earth Syst 14

Sage RF, Kubien DS (2007) The temperature response of C(3) and C(4) photosynthesis. Plant Cell Environ 30:1086–106. http://www.ncbi.nlm.nih.gov/pubmed/17661749

Smith NG, Keenan TF, Colin Prentice I, Wang H, Wright IJ, Niinemets Ü, Crous KY, Domingues TF, Guerrieri R, Yoko Ishida F, Kattge J, Kruger EL, Maire V, Rogers A, Serbin SP, Tarvainen L, Togashi HF, Townsend PA, Wang M, Weerasinghe LK, Zhou SX (2019) Global photosynthetic capacity is optimized to the environment. Ecol Lett 22

Sperry JS, Love DM (2015) What plant hydraulics can tell us about responses to climate-change droughts. New Phytologist 207:14–27. https://onlinelibrary.wiley.com/doi/10.1111/nph.13354

Sperry JS, Venturas MD, Anderegg WRL, Mencuccini M, Mackay DS, Wang Y, Love DM (2017) Predicting stomatal responses to the environment from the optimization of photosynthetic gain and hydraulic cost. Plant Cell Environ 40:816–830.

Tamm CO (1991) Nitrogen in terrestrial ecosystems: questions of productivity, vegetational changes, and ecosystem stability. Springer Science & Business Media, Berlin.

Tarvainen L, Lutz M, Räntfors M, Näsholm T, Wallin G (2016) Increased needle nitrogen contents did not improve shoot photosynthetic performance of mature nitrogen-poor scots pine trees. Front Plant Sci 7

Tarvainen L, Lutz M, Räntfors M, Näsholm T, Wallin G (2018) Temperature responses of photosynthetic capacity parameters were not affected by foliar nitrogen content in mature Pinus sylvestris. Physiol Plant 162:370–378.

Tian X, Minunno F, Schiestl-Aalto P, Chi J, Zhao P, Peichl M, Marshall J, Näsholm T, Lim H, Peltoniemi M, Linder S, Mäkelä A (2021) Disaggregating the effects of nitrogen addition on gross primary production in a boreal Scots pine forest. Agric For Meteorol 301–302

Tor-ngern P, Oren R, Oishi AC, Uebelherr JM, Palmroth S, Tarvainen L, Ottosson-Löfvenius M, Linder S, Domec J, Näsholm T (2017) Ecophysiological variation of transpiration of pine forests: synthesis of new and published results. Ecological Applications 27:118–133. https://esajournals.onlinelibrary.wiley.com/doi/10.1002/eap.1423

Wang H, Atkin OK, Keenan TF, Smith NG, Wright IJ, Bloomfield KJ, Kattge J, Reich PB, Prentice IC (2020) Acclimation of leaf respiration consistent with optimal photosynthetic capacity. Glob Chang Biol 26

Wang F, Gonsamo A, Chen JM, Black TA, Zhou B (2014) Instantaneous-to-daily GPP upscaling schemes based on a coupled photosynthesis-stomatal conductance model: correcting the overestimation of GPP by directly using daily average meteorological inputs. Oecologia 176:703–714. http://link.springer.com/10.1007/s00442-014-3059-7

Wang H, Prentice IC, Keenan TF, Davis TW, Wright IJ, Cornwell WK, Evans BJ, Peng C (2017) Towards a universal model for carbon dioxide uptake by plants. Nat Plants 3:734–741.

Wolf A, Anderegg WRL, Pacala SW (2016) Optimal stomatal behavior with competition for water and risk of hydraulic impairment. Proc Natl Acad Sci U S A 113:E7222–E7230.

Wyka TP, Żytkowiak R, Oleksyn J (2016) Seasonal dynamics of nitrogen level and gas exchange in different cohorts of Scots pine needles: a conflict between nitrogen mobilization and photosynthesis? Eur J For Res 135:483–493. http://link.springer.com/10.1007/s10342-016-0947-x

Zha T, Wang K-Y, Ryyppo A, Kellomaki S (2002) Needle dark respiration in relation to within-crown position in Scots pine trees grown in long-term elevation of CO2 concentration and temperature. New Phytologist 156:33–41. http://doi.wiley.com/10.1046/j.1469-8137.2002.00488.x

Zhao P, Chi J, Nilsson MB, Löfvenius MO, Högberg P, Jocher G, Lim H, Mäkelä A, Marshall J, Ratcliffe J, Tian X, Näsholm T, Lundmark T, Linder S, Peichl M (2022) Long-term nitrogen addition raises the annual carbon sink of a boreal forest to a new steady-state. Agric For Meteorol 324:109112.

