## supplementary material for "An eco-physiological model of forest photosynthesis and transpiration under combined nitrogen and water limitation"

Methods S1 Details regarding the optimization of stomatal conductance and leaf N content

In our work we propose a two-step plant optimization routine to determine optimal stomatal conductance ($g_{s}$) and leaf N content ($N_{m,f}$) values. In in the first step a long-term fitness proxy is maximized by optimizing $N_{m,f}$ and $g_{s}$. The two plant variables regulate on different time-scales, where $g_{s}$ regulates faster than $N_{m,f}$. We assume that $N_{m,f}$ is constant over a week’s time period and in the second step we determine the corresponding optimal daily $g_{s}$ value by optimizing the daily fitness proxy for each day within the week. Below we will give details about the two optimization steps and highlight our core assumptions.

#### First step: Long-term optimization

Let $G$ denote the instantaneous fitness reward, which we define as the instantaneous leaf-level carbon assimilation subtracted by the cost of maintaining transpiration and photosynthetic capacity. $G$ is a function of stomatal conductance ($g_{s}$), leaf N content ($N_{m,f}$), and the environmental variables ($\mathrm{ENV}$), these includes stand size and weather variables (see table 1). We assume that $g_{s}\left( t \right)$ and $N_{f}\left( t \right)$, regulate such that the weekly cumulative reward is maximized, i.e., finding $g_{s}\left( t \right)$ and $N_{f}\left( t \right)$ such that the integral of $G$ is maximized,

$$F\left[ g_{s},N_{f} \right]=\int_{t_{\mathrm{start}}}^{t_{\mathrm{end}}} G\left( g_{s}\left( s \right),N_{m,f}\left( s \right),\mathrm{ENV}\left( s \right) \right)ds .$$

Here, $t_{\mathrm{end}}$ and $t_{\mathrm{start}}$ are the start and end of a given week. Note, $g_{s}$, $N_{m,f}$, and $\mathrm{ENV}$ are functions of time. We make further assumptions: i) $N_{m,f}\left( t \right)$ optimizes on a weekly time scale and $g_{s}\left( t \right)$ on a sub-daily time scale. ii) $N_{f}\left( t \right)$ is approximately constant over a week’s period. iii) The day-to-day change of $\mathrm{ENV}\left( t \right)$ is neglectable compared to the within-day variation. iv) The daily integration of $G$ can be approximated by SDM-2 (Wang et al. 2014). With these assumptions we can formulate an approximation of the original optimization. Let $\bar{\mathrm{ENV}}\left( t \right)$ denote the average within-day value of $\mathrm{ENV}\left( t \right)$ over a given week, i.e., $\bar{\mathrm{ENV}}\left( t \right)$ represents the average day within this time period. Now we can approximate $F$,

|  | $F\left[ g_{s},N_{f} \right]\approx7\times\left[ 2\times G\left( g_{s}\left( t_{1} \right),N_{m,f},\bar{\mathrm{ENV}}\left( t_{1} \right) \right)D_{t_{1}}+2\times G\left( g_{s}\left( t_{2} \right),N_{m,f},\bar{\mathrm{ENV}}\left( t_{2} \right) \right)D_{t_{2}} \right].$ | Eq. S 1 |
| --- | --- | --- |

Here, $t_{1}$ and $t_{2}$ are solar times, and $D_{t_{1}}$and $D_{t_{2}}$ are the corresponding time interval lengths, see Wang et al. (2014) for details on how these are calculated. Note, the expression within the square brackets is the SDM-2 approximation of the daily integral of $G$. Using an optimization algorithm to maximize Eq. S 1, we obtain optimal plant variables $N_{m,f}$, $g_{s}\left( t_{1} \right)$, and $g_{s}\left( t_{2} \right)$, for a given week. In the second optimization step, optimal $g_{s}$ values are calculated for each day within the specific week.

To solve the optimization problem, we need to model the diurnal change of our weather variable inputs. For $T_{a}$ we use eq. 3 in Wang et al. (2014). For $I_{0}$ we use eq. 2 in Wang et al., 2014 together with a conversion rate to convert global radiation to PAR. We calculate $\mathrm{VPD}$ using calculated saturated vapor and daily vapor pressure data, $\mathrm{VP}$. Saturated vapor is calculated from $T_{a}$ (Alduchov and Eskridge 1996) and we assume that $\mathrm{VP}$ is constant during the day. We assume that $C_{a}$ and $\theta$ are constant during the day.

#### Second step: fine-tuning optimization

In the second optimization step, we maximize the daily integral of $G$,

$$F_{D}\left[ g_{s},N_{m,f} \right]=\int_{t_{start,D}}^{t_{end,D}} G\left( g_{s}\left( s \right),N_{m,f}\left( s \right),\mathrm{ENV}\left( s \right) \right)ds,$$

for each day within the specified week. Here, the subscript $D$ denotes a specific day within that week, and $t_{end,D}$ and $t_{start,D}$ the start and end of $D$. Analogue to the first step, we approximate $J_{D}$,

$$F_{D}\left[ g_{s},N_{m,f} \right]\approx2\left[ G\left( g_{s}\left( t_{1,D} \right),N_{m,f},\mathrm{ENV}\left( t_{1,D} \right) \right)D_{t_{1,D}}+G\left( g_{s}\left( t_{2,D} \right),N_{m,f},\mathrm{ENV}\left( t_{2,D} \right) \right)D_{t_{2},D} \right].$$

In accordance with assumption ii), we use the $N_{f}$ value attained from the first step, thus we only need to optimize $g_{s}\left( t_{1,D} \right)$ and $g_{s}\left( t_{2,D} \right)$. To speed-up computation, we use $g_{s}\left( t_{1} \right)$ and $g_{s}\left( t_{2} \right)$, obtained form the first step, as initial guess for the optimization algorithm, when determining $g_{s}\left( t_{1,D} \right)$ and $g_{s}\left( t_{2,D} \right)$.

Methods S2 Functions for upscaling from leaf-level to canopy-level $GPP$ and $E_{C}$

We assume that the intercepted irradiance can be estimated by Beer's law thus,

$$\frac{dI}{I}=-k dLAI\Longrightarrow\frac{I}{I_{0}}=e^{-k\times LAI}.$$

Here, $I$ is the irradiance and $k$ is the light extinction coefficient. The irradiance incident on a leaf, $I_{I}$, is calculated as $I_{I}(I)=k/\left( 1-m \right)I$, $m$ is the leaf transmittance (Landsberg and Sands 2010). Because we assume that $g_{s}\propto I_{I}$, and $J_{\max}\propto I_{I}$, we can use the following relationship,

|  | $\frac{I_{I}}{I_{I,0}}=\frac{g_{s}}{g_{s,top}}=\frac{J_{\max}}{J_{max,top}}=e^{-k\times LAI}.$ | Eq. S 2 |
| --- | --- | --- |

Now we can derive the equation for the upscaling of the canopy conductance,

$$g_{C}=1.6\int_{0}^{LAI} g_{s}dLAI=1.6g_{s}\int_{0}^{LAI} e^{-k\times LAI}dLAI=1.6g_{s}\frac{1-\exp\left( -k\times LAI \right)}{k}.$$

For the upscaling of gross primary production, we have to make further assumptions. We assume that all other parameter and variables are constant along the vertical canopy-axis, thus we neglect the effect of temperature reduction along this axis. Let, $I_{I_{0}}=I_{I}(I_{0})$, $J_{\max}=X_{t}J_{max,season}(T_{a},N_{m,f})$ at some time $t$, and $c_{i,\mathrm{top}}$ denotes the value of $c_{i}$ which solves Eq. 4 for the triplet $\left( g_{s,},I_{I_{0}}, J_{\max} \right)$, i.e., $c_{i}\left( g_{s},I_{I_{0}}, J_{\max} \right)=c_{i}$. Using these assumptions, we can substitute Eq. S 2 into Eq. 2,

$$J=e^{-k\times LAI}\frac{\alpha I_{I,0}+J_{\max}-\sqrt{\alpha^{2}{I_{I,0}}^{2}+2\alpha I_{I}J_{\max}\left( 1-2\theta_{J} \right) +J_{\max}^{2}}}{2\theta_{J}}=e^{-k\times LAI}J,$$

thus,

|  | $A=e^{-k\times LAI}\frac{J}{4}\frac{c_{i}-\Gamma^{*}}{c_{i}+2\Gamma^{*}}.$ | Eq. S 3 |
| --- | --- | --- |

If we substitute Eq. S 2 into Eq. 3, we get,

|  | $A=e^{-k\times LAI}g\left( c_{a}-c_{i} \right).$ | Eq. S 4 |
| --- | --- | --- |

Because Eq. S 3 and Eq. S 4 needs to be balanced, this implies,

$$g_{\mathrm{top}}\left( c_{a}-c_{i} \right)=\frac{J_{\mathrm{top}}}{4}\frac{c_{i}-\Gamma^{*}}{c_{i}+2\Gamma^{*}}.$$

This equation is solved when $c_{i}=c_{i,\mathrm{top}}$. This relation holds at any point in the vertical canopy-axis thus,

$$\int_{0}^{LAI} A\left( g_{s},I_{I}, J_{\max} \right) dLAI=\int_{0}^{LAI} e^{-k\times LAI}A\left( g_{s},I_{I_{0}}, J_{\max} \right) dLAI=A\left( g_{s,},I_{I_{0}}, J_{\max} \right)\frac{1-\exp\left( -k\times LAI \right)}{k}.$$

Methods S3 Estimating the contribution of understory vegetation to ecosystem GPP

Adopting the assumption from Tian et al. (2021), both vegetation layers (tree and understory vegetation) have the same light use efficiency ($LUE$, defined as $GPP$/absorbed light).

Let $\phi$ denote the light level above canopy (above canopy photosynthetic photon flux density), $f_{\mathrm{PAR},c}$ is the fraction of $\phi$ absorbed by the canopy layer, $f_{\mathrm{PAR},g}$ is the fraction of $\phi$ absorbed by the ground vegetation layer, $GPP_{c}$ is the $GPP$ of the canopy layer, and $GPP_{g}$ is the $GPP$ of the understory layer, $GPP_{e}=GPP_{c}+GPP_{g}$ is the ecosystem $GPP$. Then,

$$\begin{matrix} GPP_{g}=LUE\times\phi\times f_{\mathrm{PAR},g}, \\ LUE=GPP_{c}/(\phi\times f_{\mathrm{PAR},c}), \\ GPP_{e}=GPP_{c}\times(f_{\mathrm{PAR},g}+f_{\mathrm{PAR},c})/f_{\mathrm{PAR},c}. \end{matrix}$$

Note, the fraction contribution of $GPP_{g}$ to $GPP_{e}$, i.e., $GPP_{g}/GPP_{e}$, is equal to
$f_{\mathrm{PAR},g}/(f_{\mathrm{PAR},g}+f_{\mathrm{PAR},c})$. Let $\zeta=(f_{\mathrm{PAR},g}+f_{\mathrm{PAR},c})/f_{\mathrm{PAR},c}$.

Let $K_{c}$ denote the light extinction coefficient for the canopy layer, $K_{g}$ is the light extinction coefficient for the understory layer, $LAI_{c}$ is the $LAI$ of the canopy layer, and $LAI_{g}$ is the $LAI$ of the understory layer. We calculate $f_{\mathrm{PAR},c}$ and $f_{\mathrm{PAR},g}$ by Lambert-Beer’s law (eq 1 in Tian et al., 2021),

$$\begin{matrix} f_{\mathrm{PAR},c}=1-e^{-K_{c}\times LAI_{c}}, \\ f_{\mathrm{PAR},g}=e^{-K_{c}\times LAI_{c}}\left( 1-e^{-K_{g}\times LAI_{g}} \right). \end{matrix}$$

From Tian et al. (2021) we have $K_{c}$ = 0.52, $K_{g}$= 0.69, $LAI_{g}$ in fertilized stand = 1.0, $LAI_{g}$ in control stand = 0.52, thus $\zeta$≈1.2 for the fertilized stand and $\zeta$≈1.13 for the control, see Table S 1 for a full list of estimated values.

Methods S4 Parameter estimation case: no shared parameters

The parameter estimates for the case when no parameters are shared are depicted in Table S 2 . The results are summarized in Fig. S 2-Fig. S 3. For the fertilized stand (F) the RMSE and MAPE was 1.09 (1.07) g C m^-2^ day^-1^ and 16.12 (13.86) %, respectively, for the estimation of $GPP$. For $E_{C}$ the corresponding summery statistics were 0.19 (0.21) mm day^-1^ and 25.53 (35.97) %, respectively. For the control stand (C) we got the following results: For $GPP$, RMSE = 0.98 (0.86) g C m^-2^ day^-1^ and MAPE = 15.08 (13.11) %. For $E_{C}$, RMSE = 0.19 (0.22) mm day^-1^ and MAPE = 25.93 (28.77) %. The values in parenthesis are the corresponding values from the model validation. The R^2^ for all datapoints (training set + validation set) was 0.73 ($GPP$) and 0.82 ($E_{C}$) for F. The corresponding values for C were 0.75 and 0.81. The summary statistics are taken from the fitting run with the highest likelihood.

### Figures

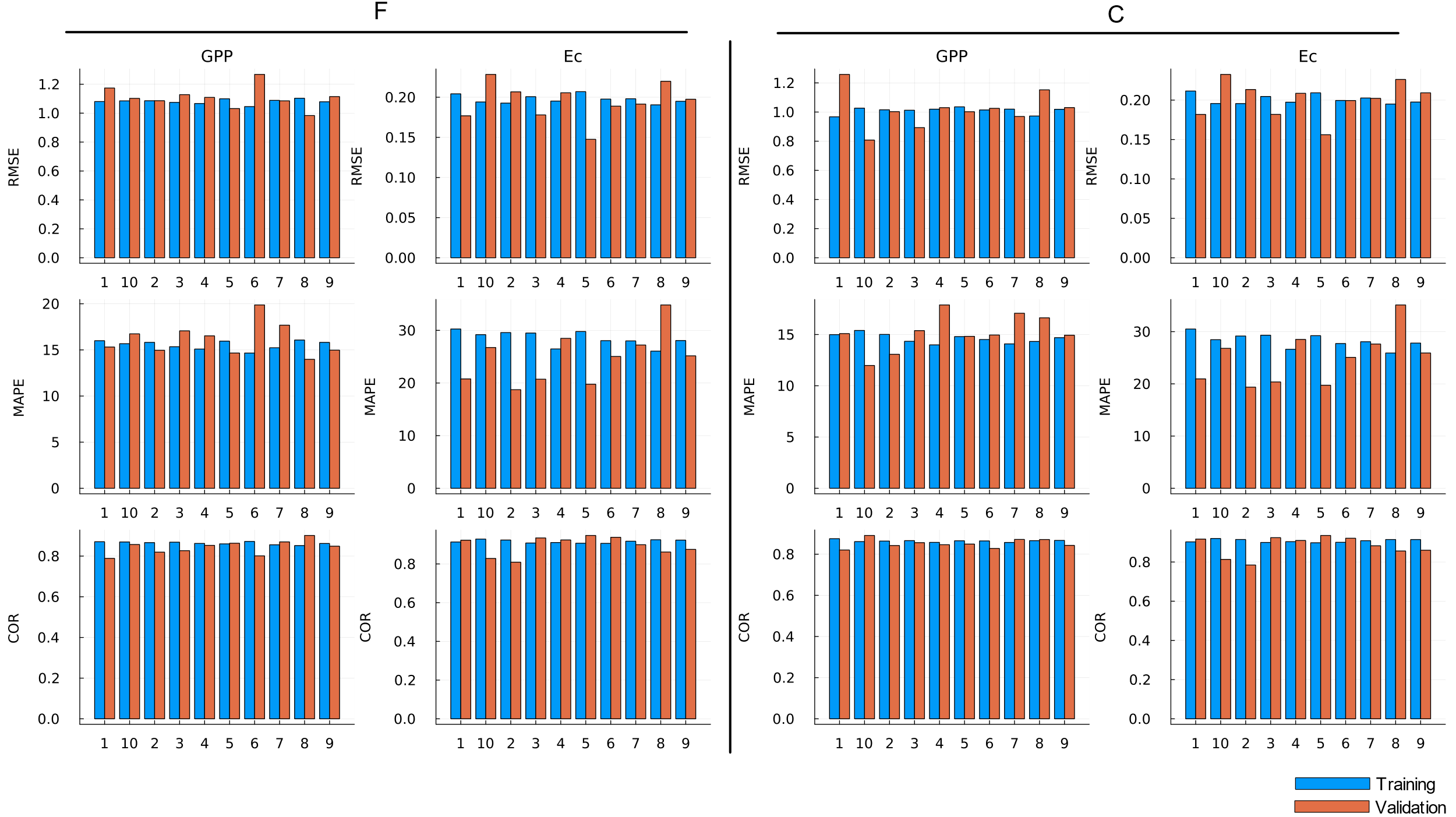

Fig. S 1 Statistical summaries of the result from parameter estimation (10 runs) (blue bars) and model validation (orange bars) for the parameter estimation case when parameters are shared among for the fertilized stand (F), left side, and control stand (C), right side: Mean absolute percentage error (MAPE), first row. Root-mean-square error (RMSE), second row. Pearson correlation coefficient (COR), third row.

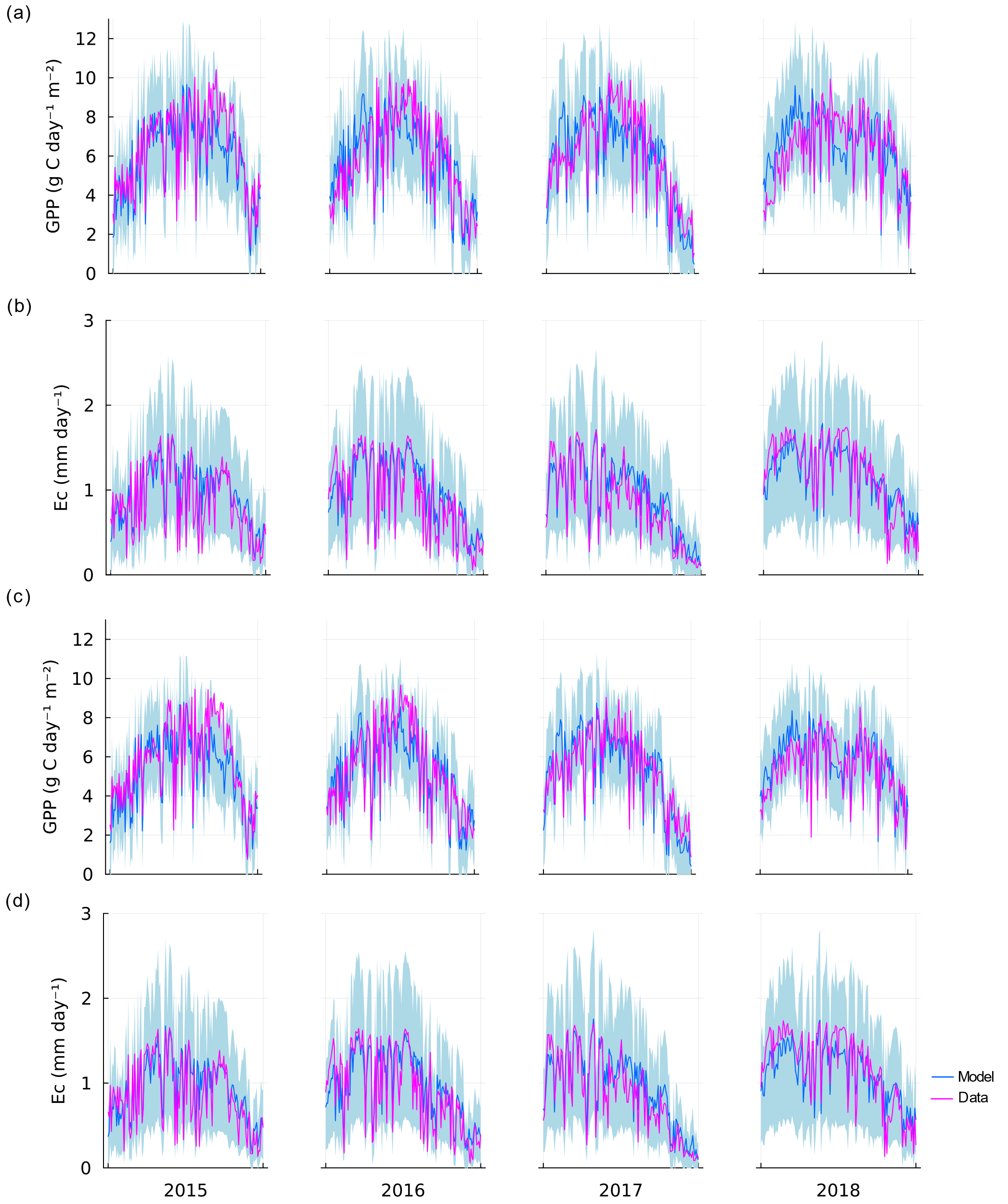

Fig. S 2 Result from the parameter estimation case when no parameters are shared between the two sand treatments, fertilized stand (a-b) and control (c-d). (a) and (c), and (b) and (d) depicts the data against the corresponding simulated estimated values for ground area gross primary production ($GPP$) and canopy transpiration ($E_{C}$), respectively. The 95% confidence interval of the fitted Laplace distribution are depicted by the light grey area.

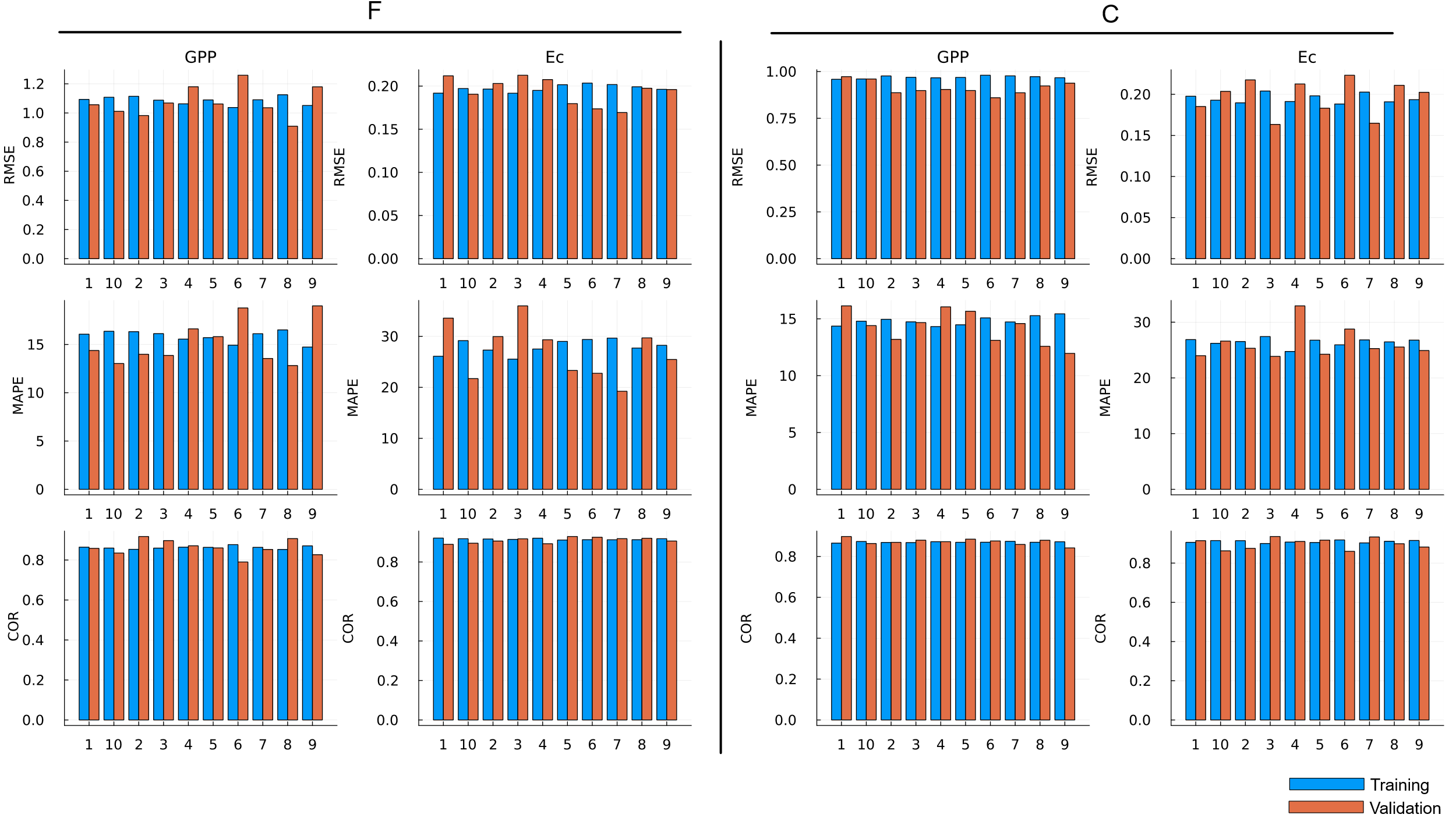

Fig. S 3 Statistical summaries of the result from parameter estimation (blue bars) and model validation (orange bars) for case when no parameters are shared among the fertilized stand (F), left side, and control stand (C), right side: Mean absolute error (MAPE), first row. Root-mean-square error (RMSE), second row. Pearson correlation coefficient (Cor), third row.

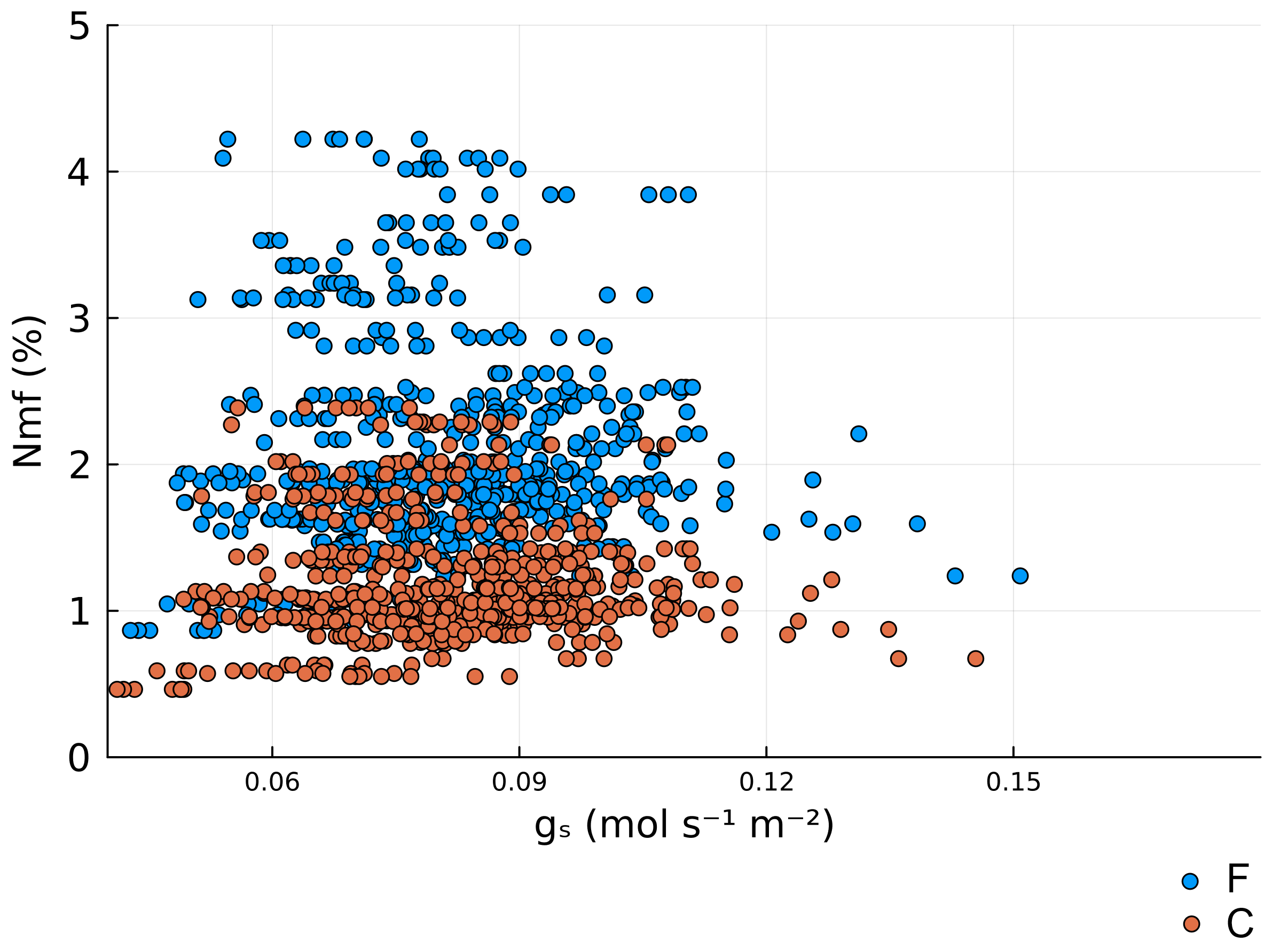

Fig. S 4 Optimal stomatal conductance ($g_{s}$) against optimal leaf mass-based N concentration ($N_{m,f}$). The blue circles correspond to the optimal trait values for the fertilized stand (F) and orange circles correspond to the control stand (C). The figure was generated by applying the model to the environmental data (Fig. 2), using parameters from Table 2 and Table 3 (treatment F and C). The correlation between $N_{m,f}$ and $g_{s}$ was -0.017 (P=.68) for F and -0.0026 (P=.95) for C.

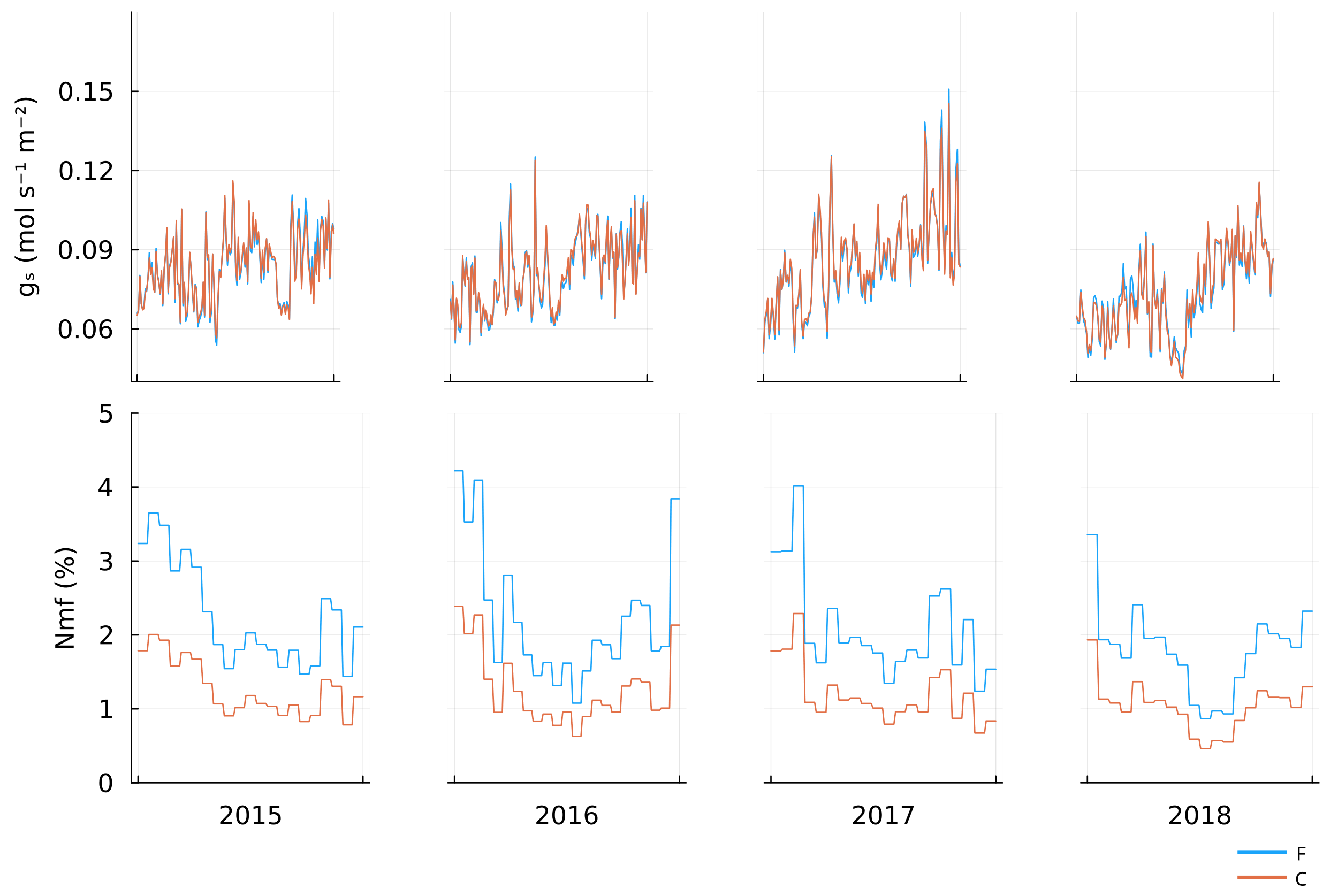

Fig. S 5 Optimal stomatal conductance ($g_{s}$), first row, and optimal leaf mass-based N concentration ($N_{m,f}$), second row, against time. The blue lines correspond to the optimal trait values for the fertilized stand (F) and orange lines correspond to the control stand (C).

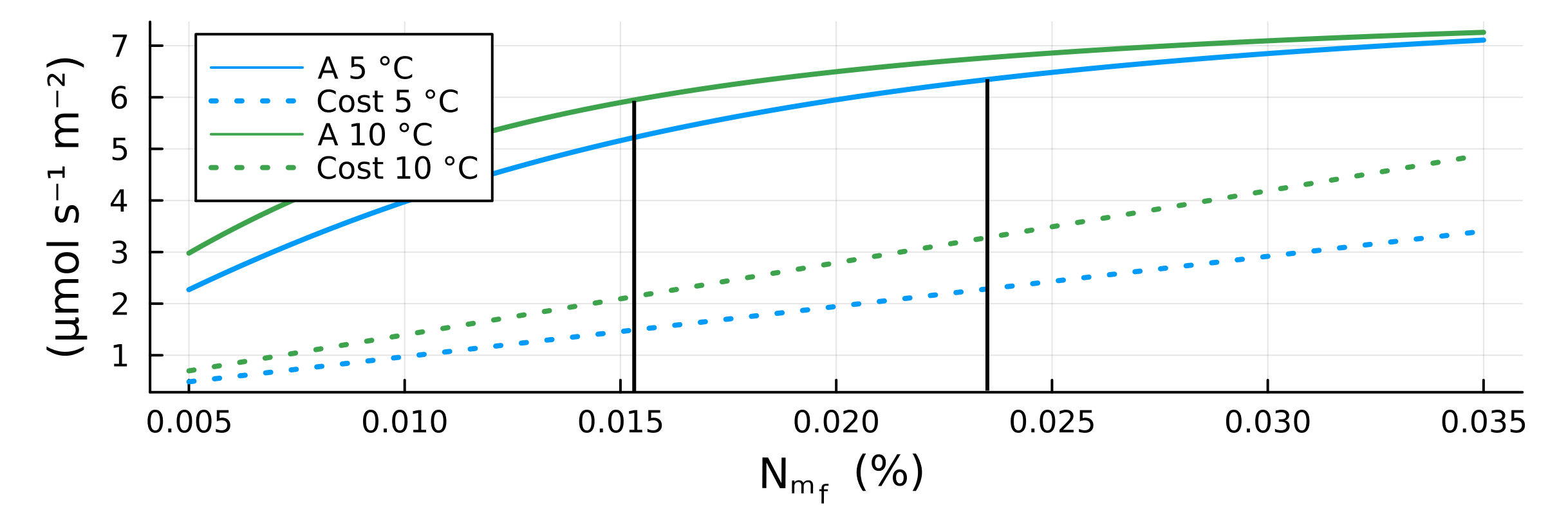

Fig. S 6 Carbon assimilation (solid curve), A, and the cost of maintaining $J_{max}$ (dotted lines), Cost, against foliage N mass-based concentration ($N_{m,f}$) for ambient temperature of 5 °C (blue line) and 10 °C (green line). Optimum $N_{m,f}$ will occur when the slope of A is equal to the slope of Cost (indicated by black horizontal lines). $g_{s}$ was set to 0.1 mol s^-1^ m^-2^ and other weather variables was set to: $I_{0}=600$ µmol s^-1^ m^-2^, $VPD=250$ Pa, and $\theta=0.175$. Parameters from Table 2 and Table 3 (Control) where used to create the figure.

### Tables

Table S 1 The fraction contribution of ground vegetation layer to eco system $GPP$ ($GPP_{g}/GPP_{e}$) and scaling factor $\zeta$ for the Fertilized stand and the control during the growth periods of 2015-2018.

| **Year** | **2015** | **2016** | **2017** | **2018** |
| --- | --- | --- | --- | --- |
| *Fertilized* | | | | |
| Canopy $LAI$ | 2.45 | 2.42 | 2.38 | 2.44 |
| $f_{\mathrm{PAR},c}$ | 0.72 | 0.72 | 0.71 | 0.72 |
| $f_{\mathrm{PAR},g}$ | 0.14 | 0.14 | 0.14 | 0.14 |
| $\zeta$ | 1.19 | 1.2 | 1.2 | 1.19 |
| $GPP_{g}/GPP_{e}$ | 0.16 | 0.17 | 0.17 | 0.16 |
| *Control* | | | | |
| Canopy $LAI$ | 2.3 | 2.28 | 2.3 | 2.21 |
| $f_{\mathrm{PAR},c}$ | 0.7 | 0.69 | 0.7 | 0.68 |
| $f_{\mathrm{PAR},g}$ | 0.09 | 0.09 | 0.09 | 0.1 |
| $\zeta$ | 1.13 | 1.13 | 1.13 | 1.14 |
| $GPP_{g}/GPP_{e}$ | 0.12 | 0.12 | 0.12 | 0.12 |

Table S 2 Estimated parameters for the case when no parameters are shared between the two stand treatments.

| **Parameter** | **Description (units)** | **Estimates** | | |
| --- | --- | --- | --- | --- |
|  |  | **F** | | **C** |
| *Trait optimization parameters* | | | | |
| $N_{u}$ | Cost parameter of maintaining $J_{\max}$ (-) | 0.0 | | 0.0045 |
| *Gross primary production model parameters* | | | | |
| $\alpha_{\mathrm{season}}$ | seasonal apex value of quantum yield peak (-) | 0.19 | | 0.17 |
| $a_{Jmax}$ | Ratio between $J_{max,opt}$ and Nitrogen concentration per leaf area ($N_{m,f}$) (mol m^-2^ s^-1^ kg leaf Kg^-1^ leaf N). | 0.01 | | 0.02 |
| $\Delta S$ | Parameter controlling when the photosynthesis is at full capacity (°C) | 18.61 | | 18.21 |
| $\tau$ | Parameter controlling the temperature delay (days) | 14.99 | | 14.97 |
| *Hydraulics model* parameters | | | | |
| $k_{sc,\max}$ | Maximum root-canopy hydraulic conductance (per leaf area) (mol m⁻² leaf s⁻¹ MPa⁻¹) | 0.00058 | 0.00065 | |
| *Variance parameters* | | | | |
| $a_{GPP,j}/b_{GPP,j}$ | The parameters which control the variance of the error distribution for GPP (g C m^-2^ ground day^-1^/-) | 0.53/0.06 | 0.59/0.03 | |

Table S 3 Pearson correlation (P-value in parenthesis) between model output (stomatal conductance $g_{s}$, optimal leaf mass-based N concentration, $N_{m,f}$, and water use efficiency, $wue$) and weather variables (radiance, $I_{0}$, mean ambient air temperature, $T_{a}$, vapor pressure deficit, $VPD$, and soil water content, $\theta$) for fertilized sand and control.

|  | Weather variables | | | |
| --- | --- | --- | --- | --- |
|  | $I_{0}$ | $T_{a}$ | $VPD$ | $\theta$ |
| *Fertilized* | | | | |
| $g_{s}$ | -0.56 (*P*<.001) | -0.43 (*P*<.001) | -0.77 (*P*<.001) | 0.32 (*P*<.001) |
| $N_{m,f}$ | 0.012 (*P*=.77) | -0.7 (*P*<.001) | -0.26 (*P*<.001) | 0.3 (*P*<.001) |
| $\mathrm{wue}$ | -0.29 (*P*<.001) | -0.47 (P<.001) | -0.57 (*P*<.001) | 0.15 (*P*<.001) |
| *Control* | | | | |
| $g_{s}$ | -0.55 (*P*<.001) | -0.39 (*P*<.001) | -0.76 (*P*<.001) | 0.38 (*P*<.001) |
| $N_{m,f}$ | 0.04 (*P*=.33) | -0.68 (*P*<.001) | -0.24 (*P*<.001) | 0.25 (*P*<.001) |
| $\mathrm{wue}$ | -0.36 (*P*<.001) | -0.48 (*P*<.001) | -0.62 (*P*<.001) | 0.084 (*P*<.001) |

### Reference

Alduchov OA, Eskridge RE (1996) Improved Magnus Form Approximation of Saturation Vapor Pressure. Journal of Applied Meteorology 35:601–609. http://publications.lib.chalmers.se/records/fulltext/245180/245180.pdf%0Ahttps://hdl.handle.net/20.500.12380/245180%0Ahttp://dx.doi.org/10.1016/j.jsames.2011.03.003%0Ahttps://doi.org/10.1016/j.gr.2017.08.001%0Ahttp://dx.doi.org/10.1016/j.precamres.2014.12

Landsberg JJ, Sands P (2010) Physiological ecology of forest production: principles, processes and models. Academic Press.

Tian X, Minunno F, Schiestl-Aalto P, Chi J, Zhao P, Peichl M, Marshall J, Näsholm T, Lim H, Peltoniemi M, Linder S, Mäkelä A (2021) Disaggregating the effects of nitrogen addition on gross primary production in a boreal Scots pine forest. Agric For Meteorol 301–302

Wang F, Gonsamo A, Chen JM, Black TA, Zhou B (2014) Instantaneous-to-daily GPP upscaling schemes based on a coupled photosynthesis-stomatal conductance model: correcting the overestimation of GPP by directly using daily average meteorological inputs. Oecologia 176:703–714. http://link.springer.com/10.1007/s00442-014-3059-7
